# Genomic characterization of a persistent, azole-resistant *C. parapsilosis* strain responsible for a hospital outbreak during the first COVID-19 wave

**DOI:** 10.1101/2024.11.11.622614

**Authors:** Michela Vumbaca, Gherard Batisti Biffignandi, Caterina Cavanna, Greta Bellinzona, Marta Corbella, Patrizia Cambieri, Johanna Rhodes, Jukka Corander, Fausto Baldanti, Davide Sassera

## Abstract

Yeasts belonging to the *Candida* genus typically reside on the mucosal surface and within the respiratory and gastrointestinal tract as commensals. Under conditions of host vulnerability, *they* can act as opportunistic pathogens, leading to various forms of candidiasis, including candidemia. Such infections can be particularly problematic when caused by isolates that exhibit resistance to antifungal drugs, which is becoming more prevalent in many regions.

One hundred and seven samples of *Candida* spp. were isolated from patients with candidemia in the hospital San Matteo in Pavia (Italy) over a period of 6 years, from 2015 to the first COVID wave in spring 2020. In order to understand the epidemiology of *Candida* infections in this hospital setting, the isolates were whole-genome sequenced which identified most as *C. parapsilosis* and *C. albicans*. Comparative genomics revealed that isolates of *C. albicans* were genomically diverse, indicating repeated introductions in the hospital from the community. *C. parapsilosis* isolates comprised two groups of highly similar isolates representing strains capable of long-term persistence in the hospital. All isolates of the main persistent group were resistant to fluconazole and presented variable levels of resistance to voriconazole and itraconazole, resulting from the Y132F substitution in erg11 and the N455D substitution in upc2. Interestingly, with the exception of the single isolate susceptible to both voriconazole and itraconazole, all the 61 isolates presented one unreported missense mutation in mrr1 (S1907C).

## INTRODUCTION

*Candida* species are one of the most common causes of fungal infections in hospital settings (1). *Candida* spp. typically colonize skin and the mucosal surface as commensal pathogens. In immunocompromised individuals they are often associated with more serious infections referred to as candidiasis which can occur as cutaneous, mucosal, or systemic infections, including those infecting multiple organs and bloodstream (this last known as candidemia) (2).

Over the past few decades the most prevalent species of *Candida* causing nosocomial infections worldwide has been *Candida albicans*, although its incidence is decreasing with a concomitant increase of non-albicans species such as *Candida auris*, *Candida glabrata*, *Candida parapsilosis*, *Candida tropicalis* and *Candida krusei* (3–5). *C. parapsilosis* has become the most common non-albicans species isolated in southern Europe, some regions of Asia and Latin America (6). This epidemiological shift has been accompanied by a worrying increase in antifungal drug resistance among *Candida* infections, including *C. parapsilosis*, mostly due to reports of resistance to the azole antifungal drug class (7). Azole resistance has been particularly reported in outbreaks, which increased during the COVID-19 pandemic, due to forced relaxing of infection control procedures and change in preventive protocols (8). Indeed the rapid spread of COVID-19 in 2020 led to a significant increase in the number of hospitalized patients with a consequent breakdown in hospital control measures. Loss of these measures, along with the use of contaminated devices and patients’ vulnerability, increased the predisposition to acquire bacterial and fungal infections, including *Candida.* It must be noted that this trend has not reverted after the end of the pandemic, possibly indicating that these azole-resistant strains are spreading rapidly (4).

Azole resistance is a complex phenomenon influenced by gene copy number variation, change in gene expression level and/or amino acid substitutions in azole targets. The primary mechanism responsible for azole resistance in *C. parapsilosis* involves mutation in the azole target gene lanosterol 14-alpha demethylase (*ERG11*), belonging to the pathway of ergosterol biosynthesis, which is necessary for fungal growth. Multiple amino acid substitutions in this gene have been reported to reduce the affinity for the drug, conferring azole resistance. Y132F in erg11 is the most common substitution detected in fluconazole-resistant strains, followed by others such as G458S, K128N, K143R or D412N (9–11). Instances of these substitutions in erg11 have been reported worldwide in *C. parapsilosis* isolates non-susceptible to fluconazole (11). Less common resistance mechanisms involve mutations in multiple genes including the transcription factors *UPC2*, *TAC1* and *MRR1*. These respectively control transcription of *ERG11* (12), of the ATP-binding cassette transporter (composed of two subunits, Cdr1 and Cdr2) (13) and of the facilitator superfamily transporter mdr1 (14). Gain-of-function mutations in *TAC1* and *MRR1* transcription factors lead to an overexpression of these two efflux pumps, enhancing the drug efflux (15, 16).

In this work we conducted a retrospective epidemiology study on *Candida* spp. isolated from patients with candidemia in the hospital IRCCS Policlinico San Matteo (Pavia, Italy) over a six year period. Different species of *Candida* were identified, most infections being caused by *C. albicans* and *C. parapsilosis*, with an increase of the latter during the first wave of the COVID-19 pandemic. Evolutionary relationships between samples were inferred through phylogenetic analysis, allowing the detection of a persistent and azole resistant *C. parapsilosis* strain in the hospital. Isolates were characterized for their antifungal resistance and genotyping analysis were conducted to investigate genotypes potentially correlated with resistance.

## MATERIALS AND METHODS

### Sample collection and species identification

Blood cultures from patients with suspected sepsis were processed using BD-Bactec FX (Becton Dickinson, Franklin Lakes, NJ, USA) employing BD BACTEC Plus Aerobic/F and Plus Anaerobic/F culture vials. When Gram staining from a positive blood culture indicated the presence of yeast, multiplex PCR (BioFire® FilmArray®) was conducted. *Candida* species grew on conventional culture and the identification of the isolates was performed using Matrix-assisted laser desorption/ionization time of flight mass spectrometry (MALDI-TOF-MS, Bruker Daltonics, Bremen, Germany) using the software Bruker Biotyper 3.1.

The 107 isolates included in the analysis represent all candidemias that occurred in the San Matteo Hospital in Pavia during the study period (January 2015 to April 2020), with the exception of the period between 2015 and 2017, when only a portion of samples isolated from affected patients could be recovered (the total number of candidemia in this period is unknown).

### Antifungal susceptibility test

Antifungal susceptibility testing was then performed using Sensititre YeastOne (SYO, Thermo Scientific Trek Diagnostic System, East Grinstead, UK). Antifungals tested were those included in the SYO panel, namely amphotericin B, anidulafungin, caspofungin, micafungin, fluconazole, itraconazole, posaconazole and voriconazole. Minimal inhibitory concentration (MIC) values obtained with SYO were interpreted according to Clinical Laboratory and Standards Institute (CLSI) species-specific breakpoints (17) or, in case of lack of these breakpoints, according to the SYO epidemiological cutoff values (ECV). Specifically, as posaconazole is not included in the CLSI guidelines, we used EUCAST v.7.0 (18) to assess resistance to this drug.

### DNA extraction, genome sequencing and genome-based species identification

For extraction of genomic DNA, cells from single colonies of each candidemia isolate were picked and grown overnight at 30°C in four ml cultures of YPD broth (1% [wt/vol] Difco yeast extract, 2% [wt/vol] Bacto peptone, 2% [wt/vol] dextrose) with shaking. DNA was then extracted with the Qiagen Genomic-tip 100/G using the Qiagen genomic buffer set (Qiagen, Hilden, Germany), following the manufacturer’s yeast protocol. Cell wall digestion was accomplished with lyticase (catalog number L2524; Sigma-Aldrich St. Louis, Missouri, USA). Library preparation and 2×150 bp paired end sequencing was performed using the Illumina NovaSeq platform.

Raw sequencing reads were quality checked and trimmed to remove low quality bases and adapter sequence using FastP v.0.23.2 (19). fastANI v.1.33 was used for genome-based species identification (20). One clinical isolate of *Candida parapsilosis* (35763_2_2) was chosen for additional long read sequencing using the MinION sequencer (Oxford Nanopore Technology, Oxford, UK).

### Genome mapping, assembly and phylogeny

*C. albicans* and *C. parapsilosis* reads of each isolates were mapped to their reference genomes, ASM18296v3 and ASM3628897v1 respectively, with Bowtie2 v.2.5.4 (21) with default settings and converted to sorted BAM format using Picard SortSam v.3.1.1 (https://broadinstitute.github.io/picard/). PCR duplicates reads were identified and marked through Picard MarkDuplicates. Genome Analysis Toolkit (GATK) v.4.4.2 HaplotypeCaller was employed for variant calling (single nucleotide polymorphism (SNPs) and indels) using default settings, while specifying samples diploidy and enabling the option -ERC GVCF (72, 22). The GVCFs were subsequently imported into a GATK GenomicsDB to merge GVCFs from multiple samples into a combined VCF (one for each *Candida* species) with the GenotypeGVCFs GATK function. Variants from the merged VCF were filtered to select high-confidence variants using the following parameters: DP <= 20, QD < 2.0, MQ < 40.0, FS > 60.0, MQRankSum < -12.5 and ReadPosRankSum < -8.0. Only SNPs were retained through GATK SelectVariants with --select-type-to-include SNP option. Samples that had genome quality (--minGenomeQuality) lower than 50 and sample depth (--minSampleDepth) lower than 30 were removed using VcfFilter v.0.2 (https://github.com/biopet/vcffilter). The multi-samples VCF resulting from the mapping procedure was converted into a FASTA alignment using ‘vcf2phylip’ (Ortiz et al. 2019) and invariant sites were removed through snp-sites v.2.5.1 (23). Pairwise SNPs distances between isolates were calculated using “Snpbreaker.R” script from P-DOR pipeline (24). Maximum-likelihood phylogenetic trees were inferred with IQTREE v.2.0.7 (25) using the MFP-GTR model and 1000 bootstrap replicates.

Illumina reads from isolates belonging to species *C. glabrata*, *C. rugosa*, *C. norvegensis* and *C. orthopsilosis* were subjected to genome assembly using Unicycler v.0.5 (26) with default parameters. The reads from the *C. parapsilosis* isolate 35763_2_2, sequenced with both Illumina and Nanopore technology were assembled into an hybrid assembly using Unicycler v.0.5 (26) with the --linear_seqs 8 option, representing the predicted presence of eight linear chromosomes in the *C. parapsilosis* reference assembly ASM3628897v1. Completeness and quality of the assembly were evaluated with BUSCO v5.8.0 (27).

### Genomic characterization of *C. parapsilosis* isolates

Genomic variants of *C. parapsilosis* were annotated with snpEff v.5.2a (28) from the VCF files, using a custom database generated from the ASM3628897v1 reference genome after annotation with Augustus (29) using the yeast mitochondrial codon tables. SnpEff was used to identify mutations present in the genes previously reported to be involved in azole resistance: *ERG11*, *FKS1*, *MRR1*, *ERG3*, *TAC1*, *UPC2* and *NDT80*. Copy number variations (CNVs) were determined from BAM files using Delly v.1.2.6. CNV calling and the output files were plotted using rd.R script (30).

## RESULTS and DISCUSSION

### *Candida parapsilosis* and *albicans* are responsible for most candidemia during the study period

Following an increased incidence of fungal candidemias observed in the IRCCS Policlinico San Matteo hospital during the first wave of the COVID-19 pandemic in Italy, we investigated the epidemiology of these infections in the hospital between January 2015 and April 2020. Our analyses include 107 isolates of *Candida* spp. isolated from 106 patients diagnosed with candidemia admitted to multiple wards. We report a notable expansion of *C. parapsilosis* and *C. albicans* between 2018 and 2020, with a spike of the latter at the beginning of the pandemic. Species identification was initially assessed through MALDI-TOF which identified 5 different species, namely *C. albicans* (*n* = 18), *C. glabrata* (now known as *Nakaseomyces glabratus* (31)) (*n* = 3), *C. norvegensis* (*n* = 1), *C. parapsilosis* (*n* = 84) and *C. rugosa* (now known as *Diutina rugosa*, (32)) (*n* = 1). Most *C. parapsilosis* isolates exhibited variable levels of resistance to azole antifungal drugs, while all isolates belonging to other species resulted to be susceptible to all tested drugs, with only a few exceptions: the three *C. glabrata* isolates show a sensible dose-dependent (S-DD) phenotype to fluconazole and itraconazole and the single *C. norvegensis* was resistant to 5-flucytosine (**Table S1**).

All isolates were subjected to Illumina whole genome sequencing, obtaining an average of 40.750.143,6262 reads per sample. Genome-based species identification was then performed, detecting three cases of discordance with the MALDI-TOF results. Specifically, three *C. parapsilosis* were re-classified as *C. orthopsilosis*, ending with a final dataset of *C. albicans* (*n* = 18), *C. glabrata* (*n* = 3), *C. norvegensis* (*n* = 1), *C. orthopsilosis* (*n* = 3), *C. parapsilosis* (*n* = 81) and *C. rugosa* (*n* = 1) (**Figure 1**). Genome-based species identification was used for subsequent analyses. The misidentification of the three *C. orthopsilosis* species as *C. parapsilosis* by MALDI-TOF is likely due to the high similarity between these closely related species. Other studies reported inconsistencies in species identification within the *C. parapsilosis* complex between MALDI-TOF results and molecular methods, suggesting that the accuracy of MALDI-TOF identification could be improved or complemented by genomic approaches (33, 34).

**Figure 1.**
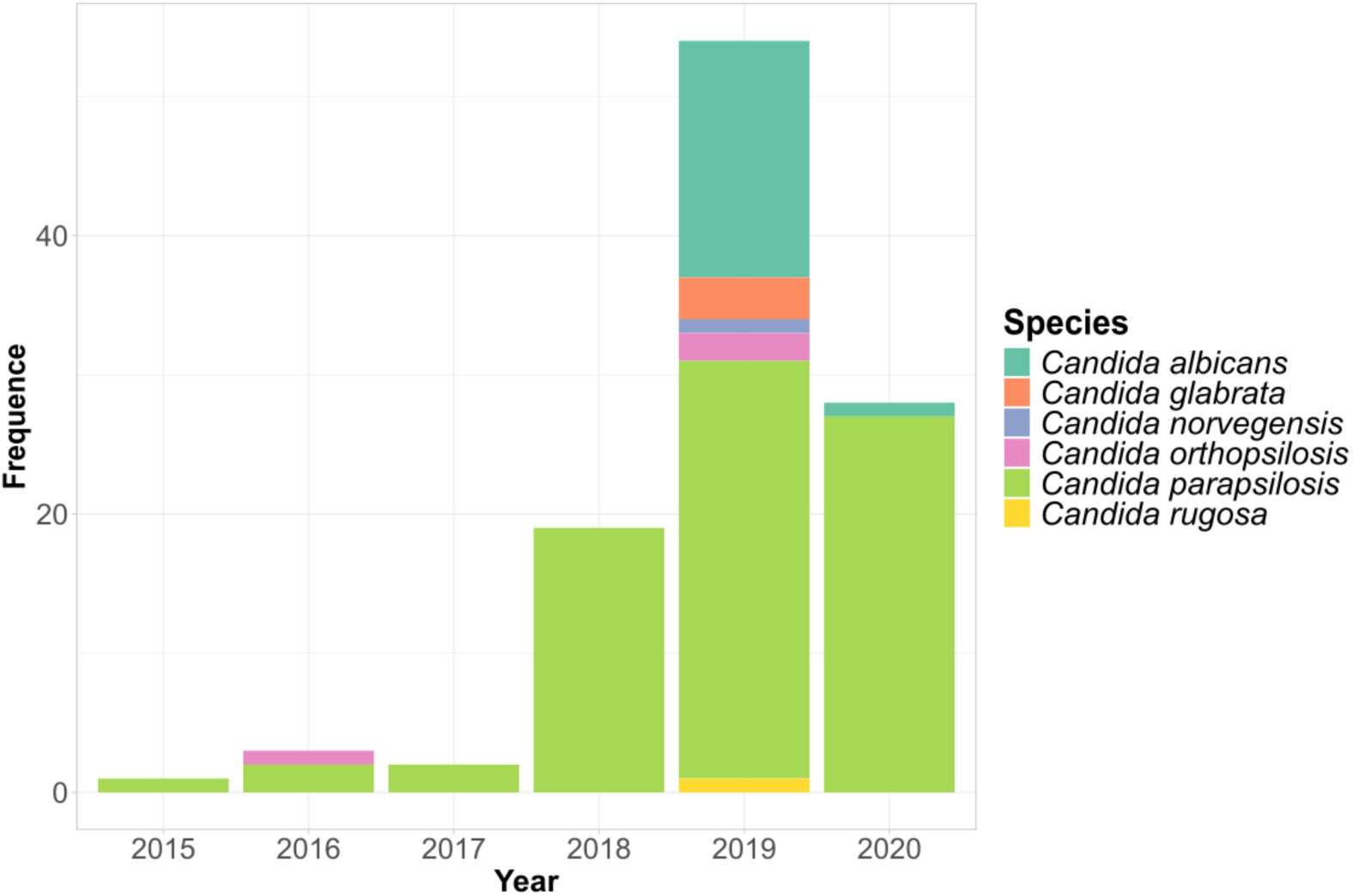
Barplot of the 107 *Candida* isolates from the hospital San Matteo in Pavia over the six years period, color coded by species. The last bar shows the isolates collected until 14 April 2020, when the sample collection for this study ended.

Genomes of isolates belonging to the four less common species were *de novo* assembled to provide references for future studies, since only a limited number of genomes are available on public databases (searching NCBI on September 2024 we found*: C. glabrata*: 69; *C. norvegensis*: 3; *C. rugosa*: 2; *C. orthopsilosis*: 7). For the two most commonly isolated species, *C. parapsilosis* and *C. albicans*, comparative genomic analyses were conducted to assess their genomic characteristics and gain insights into their spread inside San Matteo hospital. Additionally, one *C. parapsilosis* isolate was subjected to long-read sequencing and hybrid genome assembly, obtaining a chromosome level assembly of the eight chromosomes and mitochondrial genome.

### Independent introductions of *Candida albicans* into a hospital setting are not epidemiologically linked

All but one of the *C. albicans* isolates were collected in 2019 from various wards located in multiple buildings of the San Matteo Hospital (**Table S1**). To evaluate the potential epidemiological connections among the 18 *C. albicans* isolates, we performed a phylogenetic analysis, including 38 additional publicly available genomes representing the global diversity of *C. albicans*, to place the isolates from San Matteo Hospital within a broader global context (35). Information about isolates retrieved from public databases are listed in **Table S2**. Mapping to the reference allowed us to obtain a set of 41,403 whole SNP which was used to generate a maximum likelihood tree (**Figure 2**).

**Figure 2.**
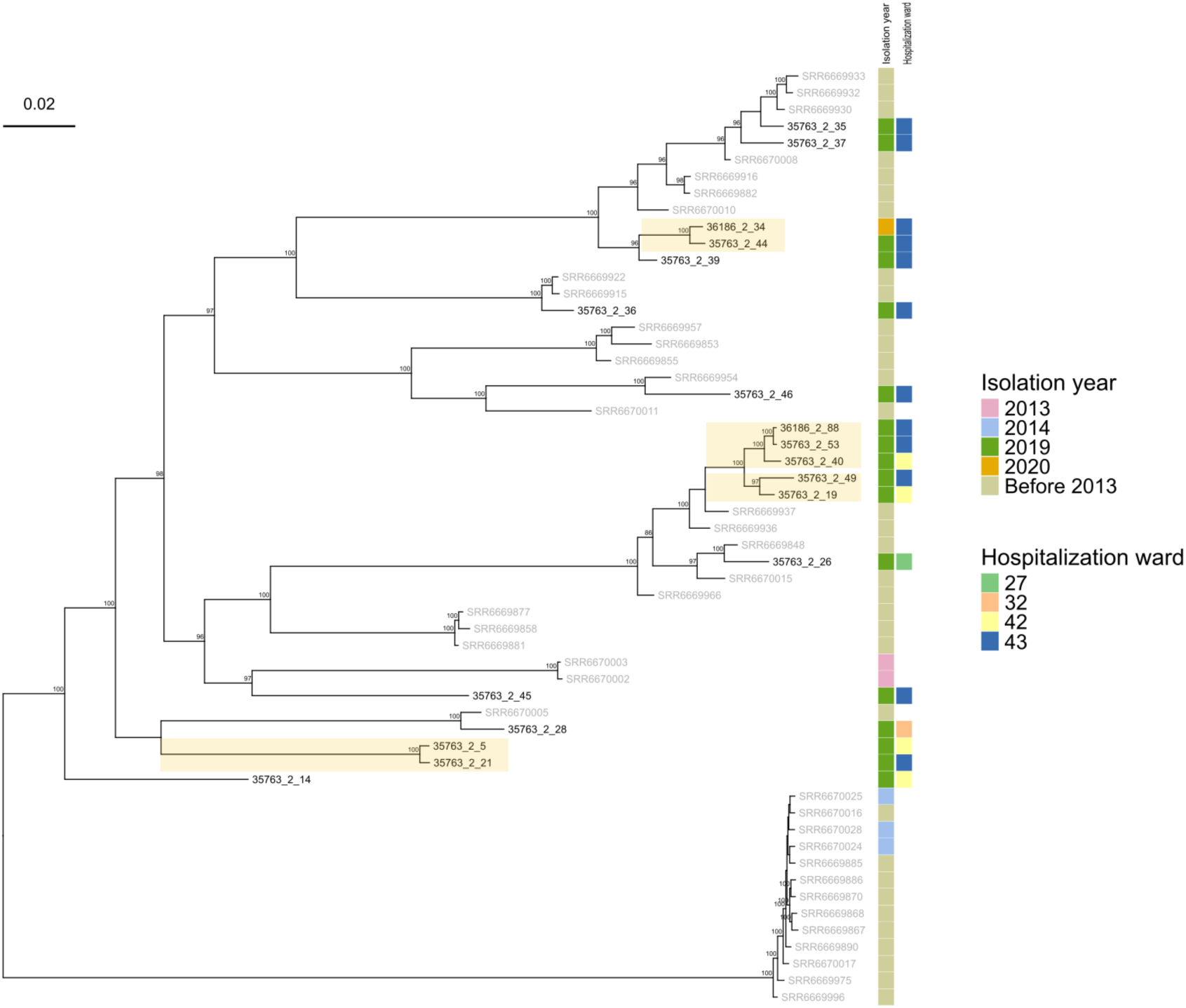
*Candida albicans* phylogeny, contextualized within a global context (**Table S2**), obtained from a whole genome SNP alignment of sequencing reads mapped to reference genome ASM18296v3, using the software IqTREE v.2.0.7. Isolate labels from San Matteo Hospital are displayed in black, while labels for publicly available genomes are shown in gray. Phylogeny is midpoint-rooted. Bootstrap values above 80 are reported on the corresponding node. For the novel isolates, year and hospital ward of isolation are indicated. The scale bar represents the branch length of the phylogenetic tree measuring the evolutionary distance between isolates.

The resulting tree revealed that the local *C. albicans* isolates are scattered on the global phylogeny, without clustering into a monophyletic group, indicating that these are independent cases that have emerged separately. However, four small monophyletic clades composed solely by genomes from the San Matteo hospital can be observed in the figure (highlighted with light yellow squares on the phylogeny), all of them composed by a limited number of genomes, from two to four (with a bootstrap support range between 97 and 100). The SNP distances within these clades are relatively high, from a minimum of 146 SNPs to a maximum of 424 SNPs (**Figure S1**), suggesting that they likely represent related lineages rather than direct epidemiological links. This distribution of evolutionary distances indicates that the analyzed isolates of this species are primarily transmitted through community-acquired infections rather than healthcare-associated outbreaks, consistent with the existing literature on *C. albicans* that reports mainly vertical transmissions (6).

The resistance profiles of *C. albicans* were determined *via* MIC values and CLSI breakpoints. None of the isolates was found to exhibit resistance to any of the antifungal agents evaluated (**Table S1**). This result is consistent with what is known about the epidemiology of this species, as while *C. albicans* infections exhibit variable levels of antifungal resistance, isolates of this species from candidemia patients generally show the lowest incidence of azole resistance among all *Candida* species (36, 37). Indeed, despite the predominance of *C. albicans* in immunocompromised patients, the emergence of antifungal resistance is more linked to non-albicans *Candida* species (38).

### *Candida parapsilosis* phylogenomics reveals introduction of a persistent strain resulting in a nosocomial outbreak

A phylogenomic approach was applied to the 81 *C. parapsilosis* genomes obtained from samples collected from various wards of the hospital between 2015 and 2020, with the addition of 34 genomes from other studies (7, 39–50), to provide broader context and a global comparison (**Figure 3**). Information about isolates retrieved from public databases are listed in **Table S2**.

**Figure 3.**
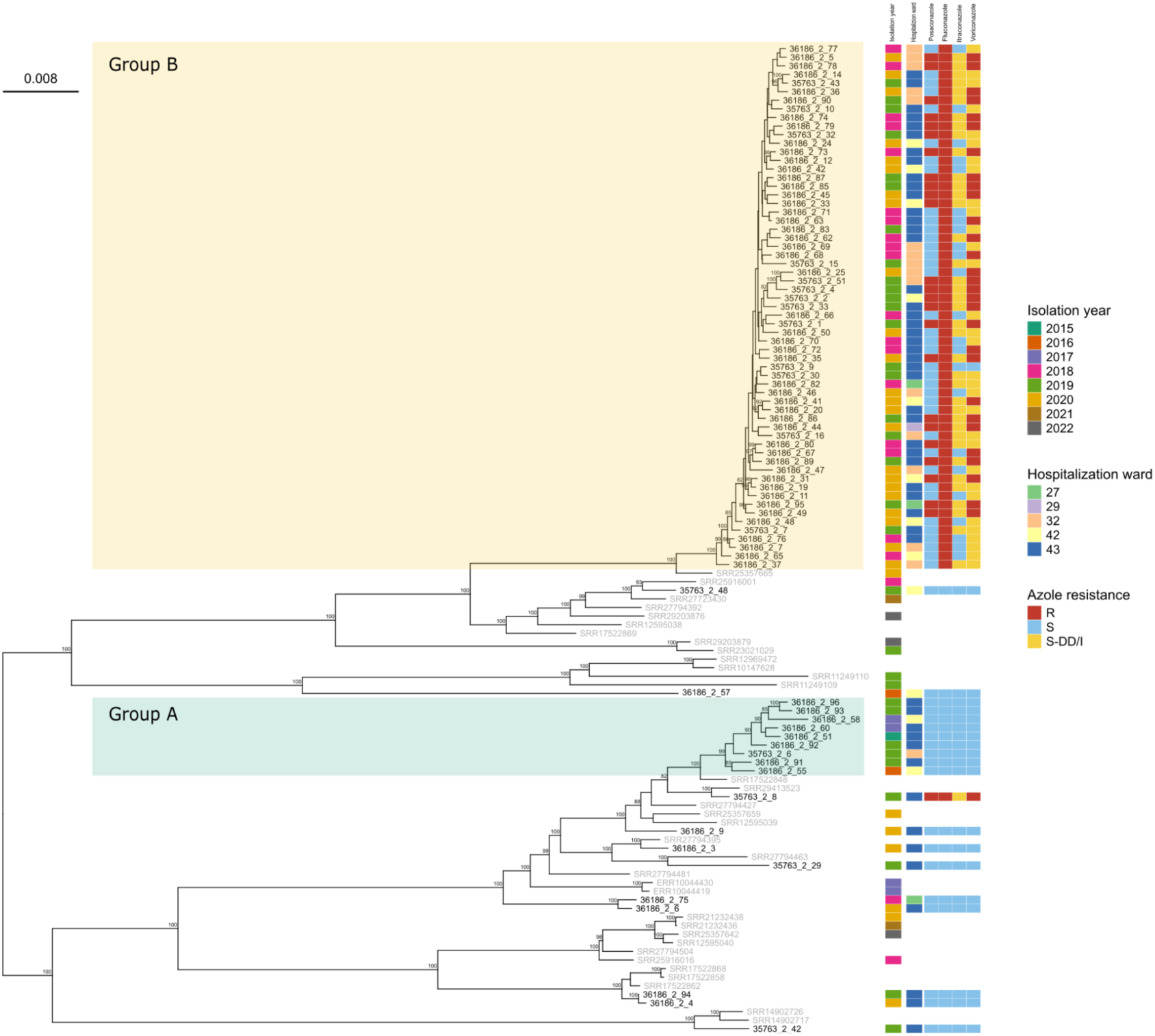
Whole genome SNP maximum-likelihood phylogeny of *Candida parapsilosis* isolates contextualized within a global context (**Table S2**). Isolate labels from San Matteo Hospital are displayed in black, while labels for publicly available genomes are shown in gray. Phylogeny is midpoint rooted. Bootstrap values above 80 are reported on the corresponding node. For novel isolates, year, hospital ward of isolation and azole resistant phenotype are indicated. The scale bar represents the branch length of the phylogenetic tree measuring the evolutionary distance between isolates.

Eleven of the novel *C. parapsilosis* isolates are scattered across the global phylogeny and do not exhibit any clear clustering either among San Matteo genomes or with any of the reference isolates in the database (**Figure 3**). This distribution suggests that, similar to what we observed with *C. albicans*, these genomes likely represent isolated cases of *C. parapsilosis* candidemia. Rather than forming a distinct clonal group, these isolates appear to have emerged separately, suggesting multiple independent introductions into the hospital over time.

The phylogeny also shows a monophyletic clade comprising nine isolates of *C. parapsilosis*, collected from 2016 to 2019 in various hospital wards (43, 42 and 32) (**Figure 3**, Group A). The average SNP distance among these isolates ranged from 3 to 60 SNPs, indicating a clear direct epidemiological link. Furthermore, phylogenomic analysis identified another monophyletic clade of 61 isolates from the studied hospital also exhibiting near-clonal similarity (**Figure 3**, Group B) supported by a bootstrap value of 100 in the phylogenetic tree. The connection among the 61 isolates was further confirmed by analyzing the pairwise SNP distances, which present differences lower than 50 SNPs among each pair (**Figure S2**).

A SNPs difference threshold to define direct epidemiological links between *C. parapsilosis* isolates has not been determined yet (40). However, multiple studies identified low genetic variation among isolates belonging to outbreaks (e.g. below 20 SNPs in (51)). In the case of the two monophyletic clades described here, the average SNP distances, slightly higher than what reported in previous outbreaks, could be attributed to the extended presence of the two strains in the hospital, both having persisted for at least three years. The two monophyletic clusters can thus be classified as two single strains capable of persisting over time, gradually accumulating a low but significant number of SNPs. Interestingly, all isolates belonging to the smaller clade were sensitive to all azoles, which could explain its limited success and eventual disappearance, leading to only a small number of candidemia cases, compared to the more successful strain that resulted in 61 infections, all caused by azole resistant isolates.

This more successful strain likely entered San Matteo hospital in 2018 and there it thrived, probably aided by its resistance to fluconazole. Presence of this resistance is relatively common in nosocomial *C. parapsilosis*, reported at rates between 5 and 20%, with increases up to 90% in outbreaks, and has been linked to persistence and spread of strains in the hospital environment (11, 52). During the following three years, this strain infected multiple patients and developed resistance to voriconazole and itraconazole as well. In 2020, during the first wave of COVID-19, it caused a bona fide outbreak, with 27 blood infections in the span of 95 days. A similar rise in candidemia incidence during the pandemic has been reported in various countries, as a result of the modifications to infection prevention protocols which allowed less common species like *C. parapsilosis* to proliferate, resulting in an accelerated rate of hospital-acquired infections (53–55). Due to the impact of this strain and its interesting pattern of drug resistance, we selected one isolate for long read sequencing with Oxford Nanopore technology, allowing us to obtain a chromosome-level assembly (BUSCO score: C:97.3% [S:97.0%, D:0.3%], F:0.3%, M:2.4%). This novel genome, consistent both for genome length (13.0 Mbp to 13.1 Mbp) and number of genes (5927 to 5958) with the ASM3628897v1 reference genome of this species, is here provided as a resource for future studies on genomic epidemiology of *C. parapsilosis* and on evolution of resistance to azoles in this species.

### *Candida parapsilosis* determinants of resistance to azoles

Out of 81 *C. parapsilosis* isolates, the 61 belonging to the main persistent strain showed variable profiles of resistance to azoles. All 61 isolates resulted resistant to fluconazole with MICs ranging from 16 to 256 μg/mL, while MICs to itraconazole, posaconazole and voriconazole were even more variable, resulting in all combinations of susceptible, intermediate or resistant phenotypes against these three last azoles (**Table S1**). The remaining 20 *C. parapsilosis* isolates, comprising the second, less persistent strain, were susceptible to all tested azoles, with the single exception of 35763_2_8 which displayed elevated MICs to fluconazole, posaconazole and voriconazole and intermediate MIC to itraconazole (256 μg/mL for fluconazole, 0.125 μg/mL for posaconazole, 0.1 μg/mL for voriconazole and 0.25 μg/mL to itraconazole). All the 81 isolates displayed susceptibility to echinocandins (micafungin, caspofungin and anidulafungin, MIC: 0.25-2 μg/mL, amphotericin B (MICs ≤ 1 μg/mL) and 5-Flucytosine (MICs ≤ 2 μg/mL) with the single exception of isolate 36186_2_91 which resulted to have intermediate MIC to anidulafungin and micafungin.

Considering the pattern of resistance to azoles, the genomes of all 81 isolates of *C. parapsilosis* were analyzed looking for genetic determinants of resistance. The presence of non-synonymous substitutions were investigated in specific genes: *ERG11*, *FKS1*, *ERG3*, *MRR1*, *UPC2*, *TAC1* and *NDT80* (**Table 1**), which have previously reported to be involved in resistance, also considering that azole resistance can be conferred by multiple mutations in different genes associated with the same pathways possibly evolving in parallel (15, 12, 38, 56). All the 61 fluconazole-resistant isolates belonging to the persistent strain harbor the well known Y132F substitution in erg11, which is absent in all susceptible isolates. Mutations in this gene, responsible for ergosterol production, are the primary cause of fluconazole resistance, not only in *C. parapsilosis* but also in *C. albicans*, *C. tropicalis* and *C. auris* (57–59). The Y132F substitution is the most common mutation in erg11 in *C. parapsilosis,* and has been well described as having a role in the clonal spread of fluconazole-resistance strains in hospital outbreaks (52, 60, 61). None of the other common mutations in erg11, G458S, K128N, and K143R (61, 62) were detected in our dataset.

**Table 1.** Pattern of mutations in genes responsible for azole resistance in the isolates of the main persistent strain of isolates. R: resistance; S: susceptible, I: intermediate, fluc: fluconazole, voric: voriconazole, itrac: itraconazole, posac: posaconazole

| Gene | Amino acid substitution in translated protein | Phenotypic Resistance (Persistent Strain) |
| --- | --- | --- |
| *ERG11* | Y132F | R fluc, S voric, itrac and posac n=1/1 |
|  |  | R fluc, I voric, S itrac and posac n=17/17 |
|  |  | R fluc and voric, S itrac and posac n=5/5 |
|  |  | R fluc and voric, I itrac and S posac n= 3/3 |
|  |  | R fluco, I itrac and voric and S posac n=11/11 |
|  |  | R fluc and posac, I itrac and voric n=3/3 |
|  |  | R fluc, voric and posac and I itrac n= 21/21 |
| *MRR1* | S1907C | R fluc, S voric, itrac and posac n=0/1 |
|  |  | R fluc, I voric, S itrac and posac n=17/17 |
|  |  | R fluc and voric, S itrac and posac n=5/5 |
|  |  | R fluc and voric, I itrac and S posac n= 3/3 |
|  |  | R fluco, I itrac and voric and S posac n=11/11 |
|  |  | R fluc and posac, I itrac and voric n=3/3 |
|  |  | R fluc, voric and posac and I itrac n= 21/21 |
| *UPC2* | N455D | R fluc, S voric, itrac and posac n=1/1 |
|  |  | R fluc, I voric, S itrac and posac n=17/17 |
|  |  | R fluc and voric, S itrac and posac n=5/5 |
|  |  | R fluc and voric, I itrac and S posac n= 3/3 |
|  |  | R fluco, I itrac and voric and S posac n=11/11 |
|  |  | R fluc and posac, I itrac and voric n=3/3 |
|  |  | R fluc, voric and posac and I itrac n= 21/21 |

Mutations in *UPC2*, which encodes for a transcription factor that regulates the ergosterol biosynthesis by modulating the *ERG11* activity, can lead to an increased production of ergosterol allowing the organism to evade the effects of these antifungals (63). All *C. parapsilosis* isolates belonging to the main persistent group (*n* = 61) harbor a non-synonymous SNP (nsSNP) in this gene, N455D. Although *UPC2* is a well-studied gene due to its involvement in resistance mechanisms, only few studies have reported this mutation (7, 64), specifically indicating its involvement resistance to the imidazole drug ketoconazole (64). A comprehensive molecular characterization of this gene and its mutants could improve our understanding of the mechanisms underlying resistance to a broader range of azoles, beyond the classical ones.

We also detected a previously unreported nsSNP substitution in *MRR1*, a zinc cluster transcription factor which influences the expression of Cdr1 and Mdr1 efflux pumps, with the result of an increased secretion of azole from the cell (65). Mutations in this gene, such as G1810A, have been reported to lead to resistance to fluconazole and voriconazole (15). The novel nsSNP causes a serine to cysteine (S1907C) amino acid substitution, and is present in all 61 persistent group isolates, with the single exception of 35763_2_9, which interestingly is the only isolate of this strain susceptible to the other three antifungal drugs: posaconazole, itraconazole and voriconazole. The absence of this mutation in this single isolate supports its potential implication in the development of resistance to itraconazole and voriconazole. Finally, analyses of *ERG3*, *TAC1* and *NDT80* genes, also reported to be involved in development of azole resistance (12, 16, 66), showed no mutation in any isolate.

All isolates displayed susceptible echinocandin MICs, with the exception of 36186_2_91 which displayed intermediate MICs for anidulafungin and micafungin. We therefore assessed *FKS1* for mutations that could explain this observation, as catalytic subunits of fks1 are the target of echinocandins drugs which lead to cell death, disrupting the cell wall (67). Through the emergence of mutations in this gene, *Candida* spp. can evade the effects of these antifungal drugs, and develop an echinocandin-resistant phenotype (12, 15). Analysis of *FKS1* coding regions showed no mutations, as expected considering that all the isolates of our dataset are susceptible to these antifungal drugs. Isolate 35763_2_7 harbored a glycine to serine substitution (G268S) within fks1, this mutation is not reported in the literature.

Chromosomes aneuploidy is one of the mechanisms through which organisms can adapt to environmental changes, providing a selective advantage during clinical treatment (68). Copy number variations (CNVs) were thus investigated, as they have been reported to drive azole resistance in *C. parapsilosis* as in other *Candida* species (69, 70). The chromosome names provided here adhere to the nomenclature used in the reference genome ASM3628897v1. Variations in chromosome ploidy were found in 16 of the 81 isolates of *C. parapsilosis*. Specifically, trisomy of different chromosomes was detected (**Figure S3**). Chromosome CP137568.1 (where *ERG11* and *TAC1* genes are present) trisomy was the most common, found in 11 isolates, all belonging to the main persistent group resistant to azoles, trait that may be linked to the increased expression of the *ERG11* and *TAC1* genes, both located on this chromosome. This result was confirmed by the read coverage of the chromosome level hybrid assembly of isolate 35763_2_2, which is one of the 11 containing this trisomy. Other chromosomal trisomies were reported in only one isolate: in 35763_2_48 for chromosome CP137570.1, in 36186_2_12 for chromosome CP137567.1 and in 36186_2_58 for chromosome CP137566.1. These chromosomes do not harbor any of the genes known to be determinants of antifungal resistance. Lastly, aneuploidy was detected in five different chromosomes (CP137570.1, CP137568.1, CP137566.1, CP137565.1, CP137569.1) in isolate 35763_2_42.

## CONCLUSION

In this work we provide insight into the epidemiology of *Candida* spp. isolated from the hospital San Matteo, Pavia, Italy over a six year period, concluding during the first wave of COVID-19 in April 2020. Our dataset shows, as previously reported (71), that the onset of the COVID-19 pandemic caused a significant rise in pathogenic infections, highlighted by a notable increase in cases of candidemia caused by *C. parapsilosis*. Within the 81 *C. parapsilosis* isolates we obtained, we were able to reveal and describe the introduction of a persistent strain, represented by 61 isolates, that resulted in an outbreak in the Italian hospital. All the persistent strain’s isolates are strongly genetically related, exhibiting a low number of SNPs that suggest a direct epidemiological connection. In terms of resistance, these isolates are all characterized by a fluconazole resistant phenotype, but interestingly, by variable levels of resistance to other azoles. Mutations in genes associated with azole resistance have been investigated, allowing us to identify, alongside known mutations, an amino acid substitution not previously reported in literature. Determining whether this mutation is linked to the variations in the resistance levels would require further ad-hoc studies and experimental validation. Our findings highlight the critical need for ongoing surveillance in hospital settings to detect outbreaks of resistant strains and enforce control measures to prevent their spread. Additionally, our newly assembled genomes, including a chromosome-level *C. parapsilosis* assembly, will be made available as a reference for future studies.

