## Supplemental material, figure and tables for "Genomic characterization of a persistent, azole-resistant *C. parapsilosis* strain responsible for a hospital outbreak during the first COVID-19 wave": Figure_S1.pdf

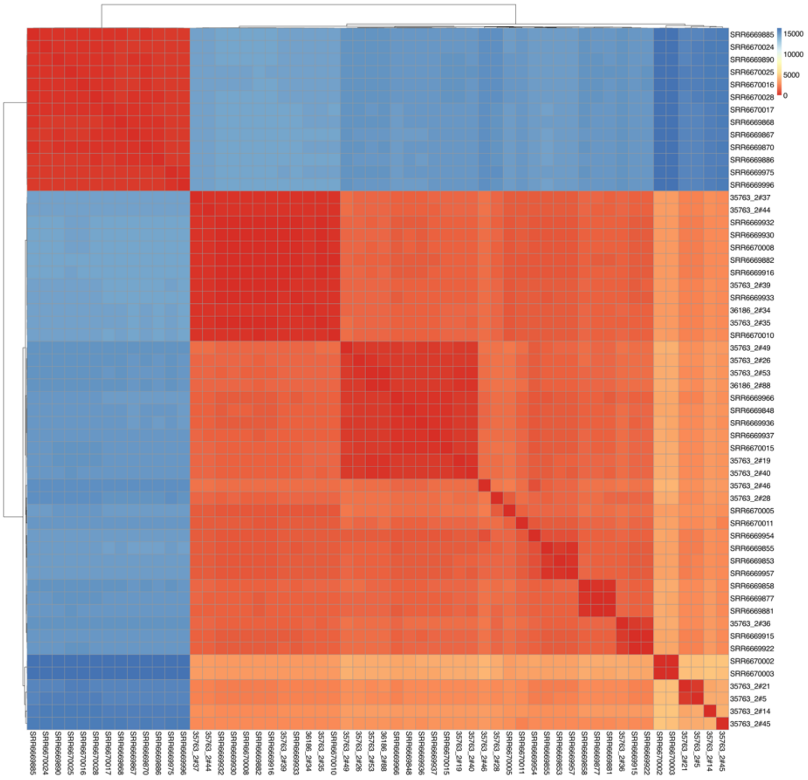


**Figure S1.** Heatmap of *C. albicans* SNPs showing the SNPs distance between all pairs of genomes included in the analysis, comprising isolates from San Matteo Hospital in Pavia and genomes retrieved from public datasets.
