## Supplemental material, figure and tables for "Genomic characterization of a persistent, azole-resistant *C. parapsilosis* strain responsible for a hospital outbreak during the first COVID-19 wave": Figure_S3.pdf

**Figure S3.** Delly plot of the 81 *C. parapsilosis* isolates representing chromosome aneuploidy.

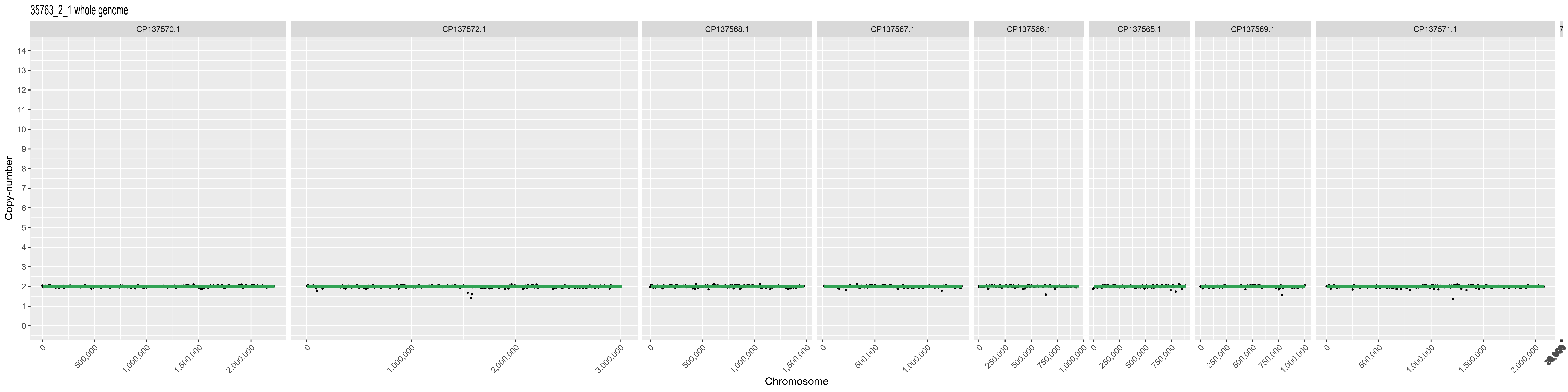

35763_2_1 whole genome
CP137570.1
CP137572.1
CP137568.1
CP137567.1
CP137566.1
CP137565.1
CP137569.1
CP137571.1
Copy-number
Chromosome

35763_2_2 whole genome

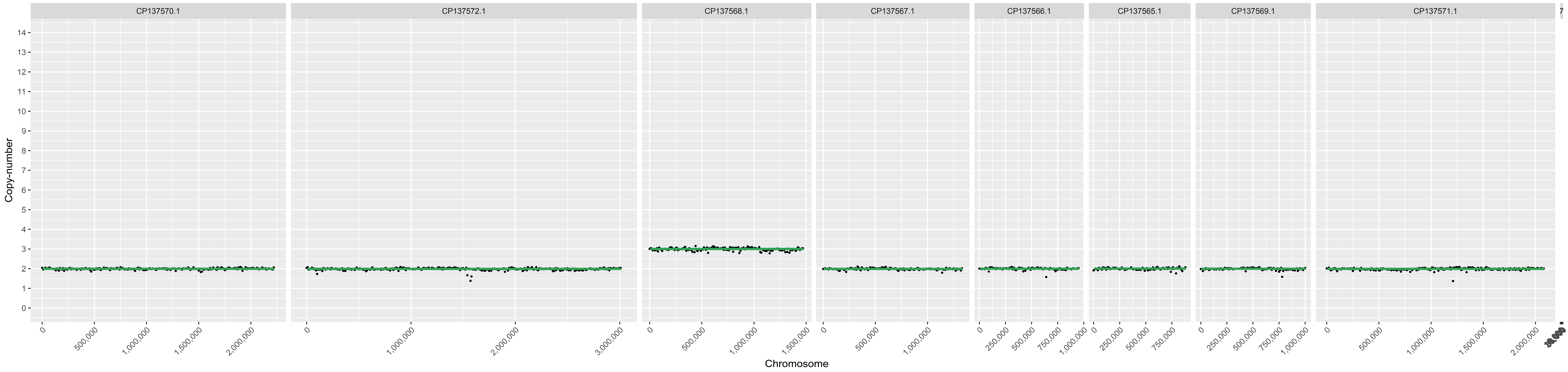

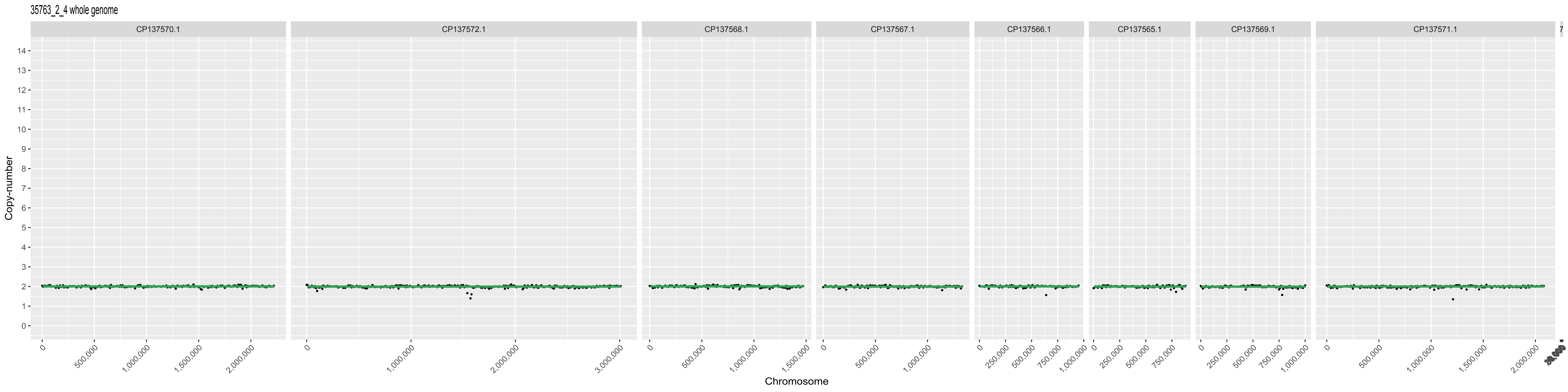

35763_2_4 whole genome
CP137570.1
CP137572.1
CP137568.1
CP137567.1
CP137566.1
CP137565.1
CP137569.1
CP137571.1
Copy-number
Chromosome

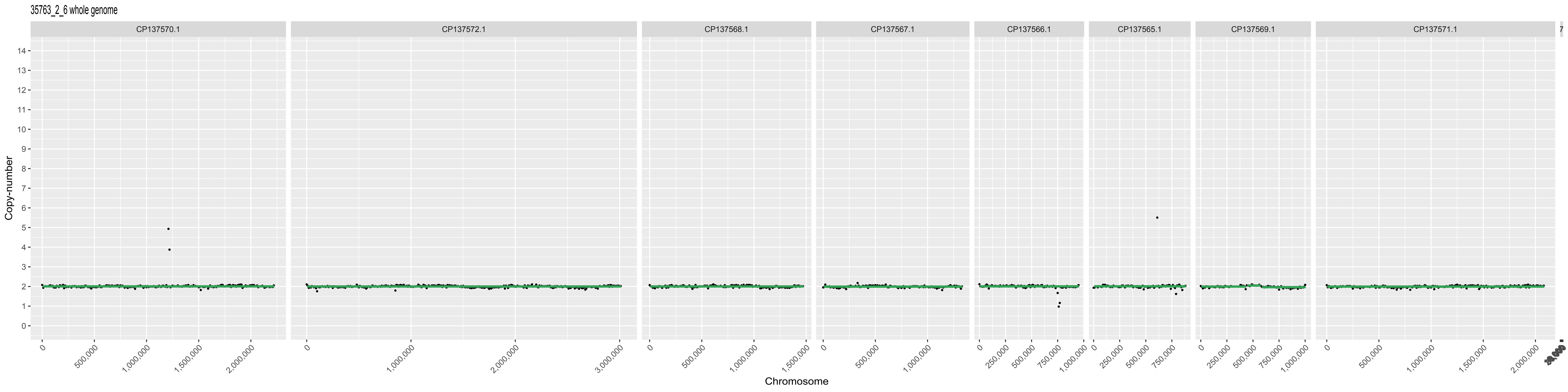
35763_2_6 whole genome
CP137570.1
CP137572.1
CP137568.1
CP137567.1
CP137566.1
CP137565.1
CP137569.1
CP137571.1
Copy-number
Chromosome

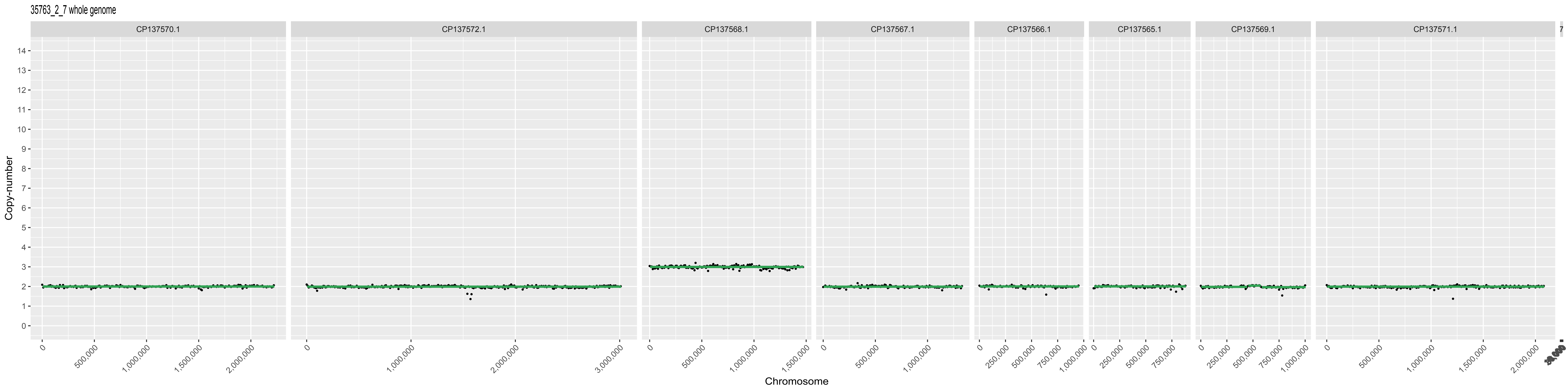
35763_2_7 whole genome
CP137570.1
CP137572.1
CP137568.1
CP137567.1
CP137566.1
CP137565.1
CP137569.1
CP137571.1
Copy-number
Chromosome
0
500,000
1,000,000
1,500,000
2,000,000
3,000,000
250,000
750,000

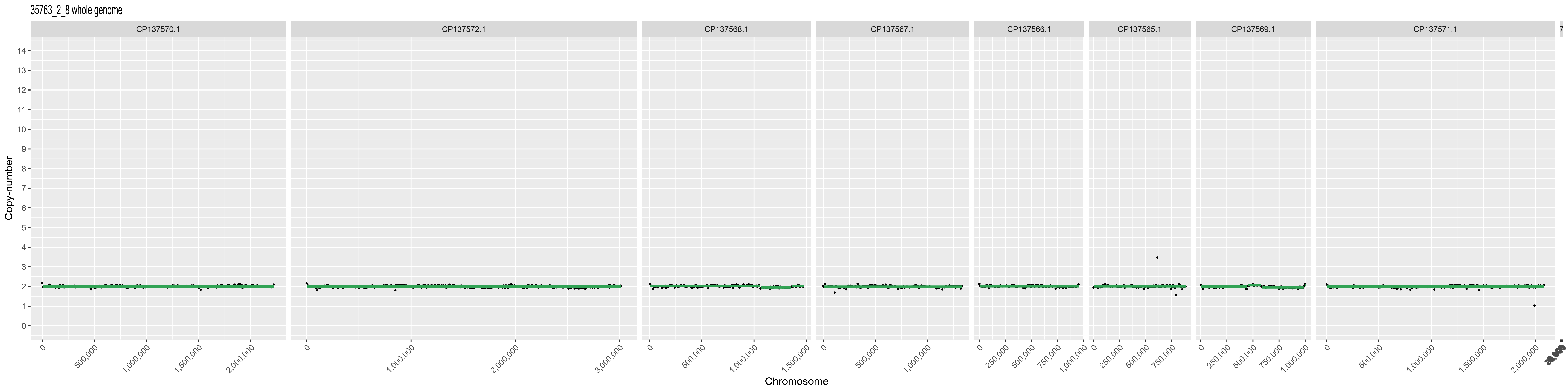

35763_2_8 whole genome
CP137570.1
CP137572.1
CP137568.1
CP137567.1
CP137566.1
CP137565.1
CP137569.1
CP137571.1
Copy-number
Chromosome

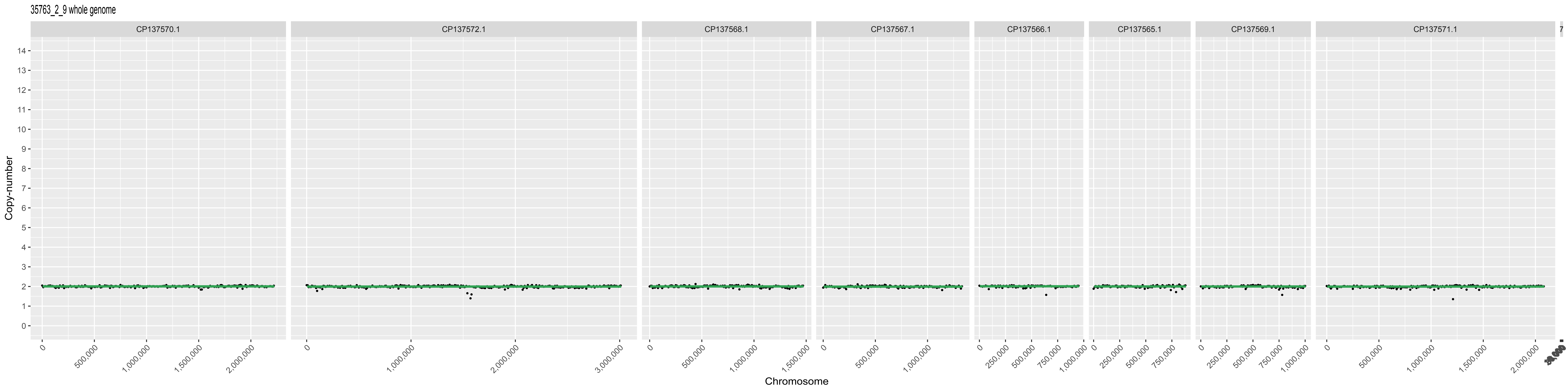
35763_2_9 whole genome
CP137570.1
CP137572.1
CP137568.1
CP137567.1
CP137566.1
CP137565.1
CP137569.1
CP137571.1
Copy-number
Chromosome

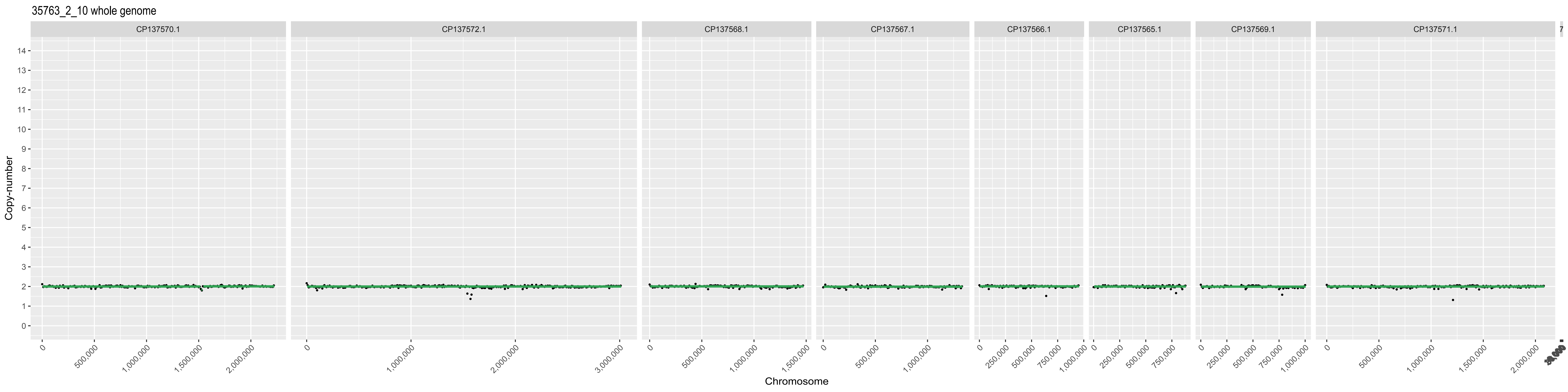

35763_2_10 whole genome
CP137570.1
CP137572.1
CP137568.1
CP137567.1
CP137566.1
CP137565.1
CP137569.1
CP137571.1
Copy-number
Chromosome

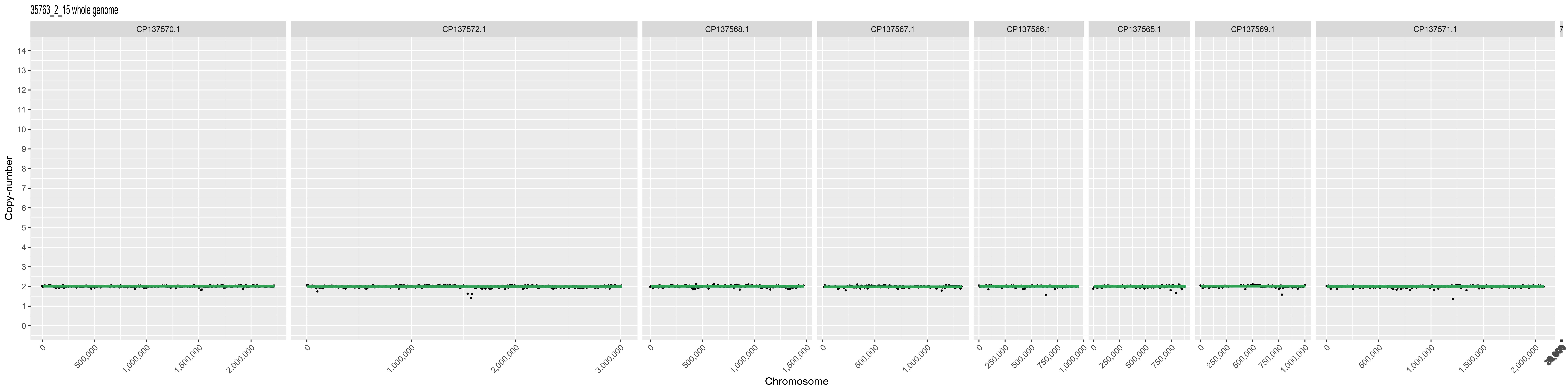
35763_2_15 whole genome
CP137570.1
CP137572.1
CP137568.1
CP137567.1
CP137566.1
CP137565.1
CP137569.1
CP137571.1
Copy-number
Chromosome
0
500,000
1,000,000
1,500,000
2,000,000
3,000,000
250,000
750,000

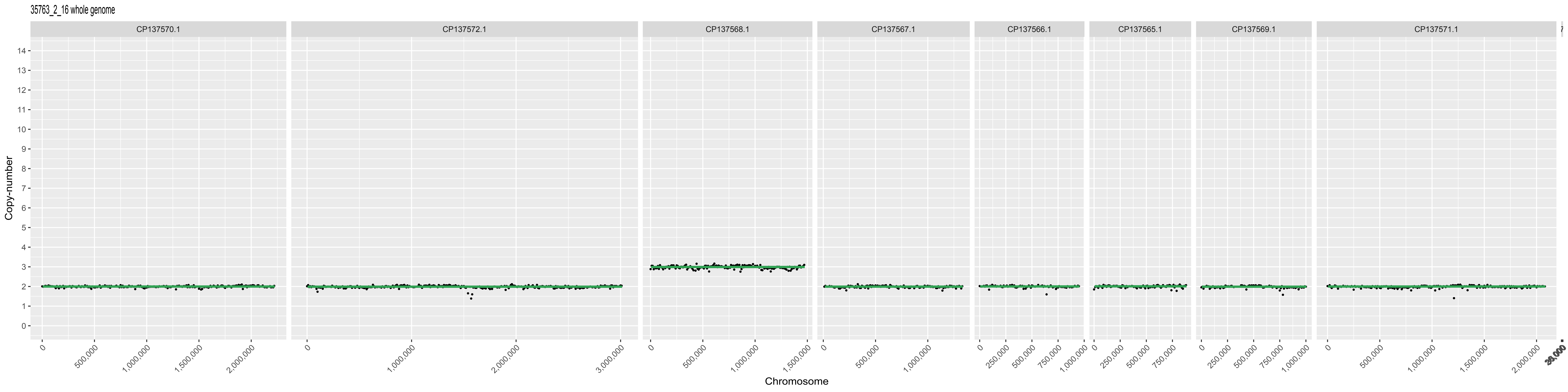
35763_2_16 whole genome
CP137570.1
CP137572.1
CP137568.1
CP137567.1
CP137566.1
CP137565.1
CP137569.1
CP137571.1
Copy-number
Chromosome
0
500,000
1,000,000
1,500,000
2,000,000
3,000,000
250,000
750,000

35763_2_29 whole genome

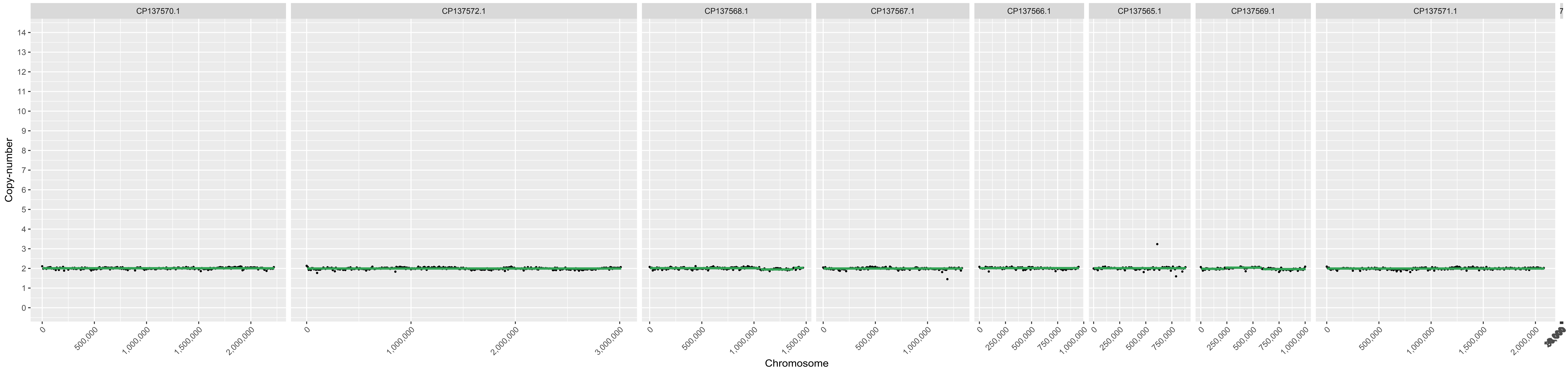

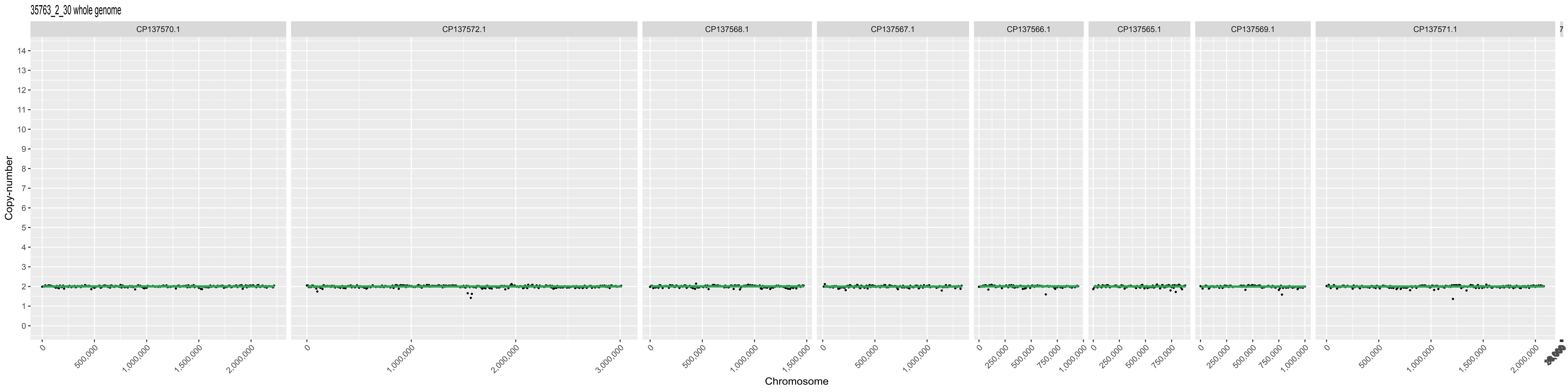
35763_2_30 whole genome
CP137570.1
CP137572.1
CP137568.1
CP137567.1
CP137566.1
CP137565.1
CP137569.1
CP137571.1
Copy-number
Chromosome

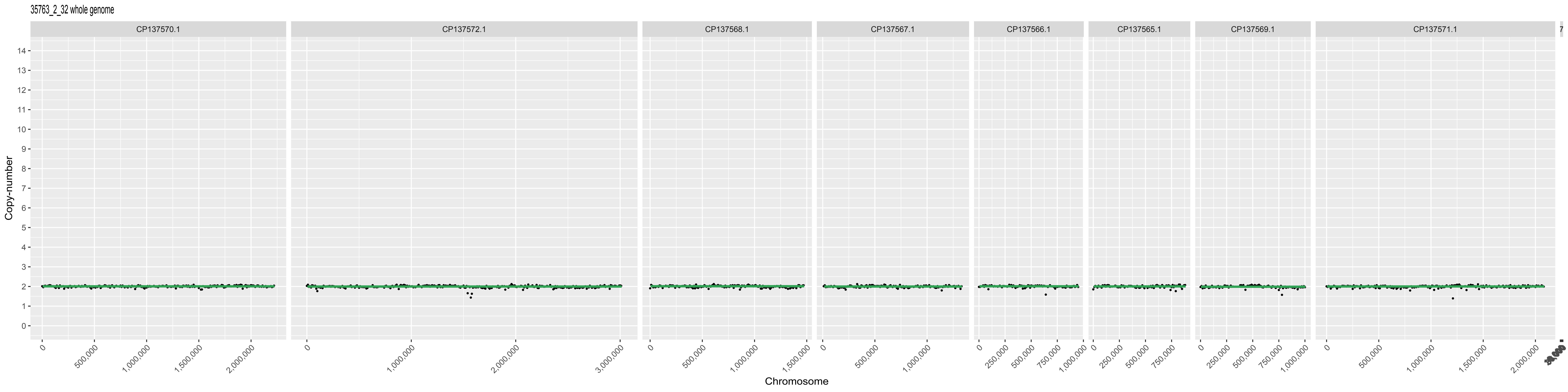
35763_2_32 whole genome
CP137570.1
CP137572.1
CP137568.1
CP137567.1
CP137566.1
CP137565.1
CP137569.1
CP137571.1
Copy-number
Chromosome
500,000
1,000,000
1,500,000
2,000,000
3,000,000
250,000
750,000

35763_2_33 whole genome

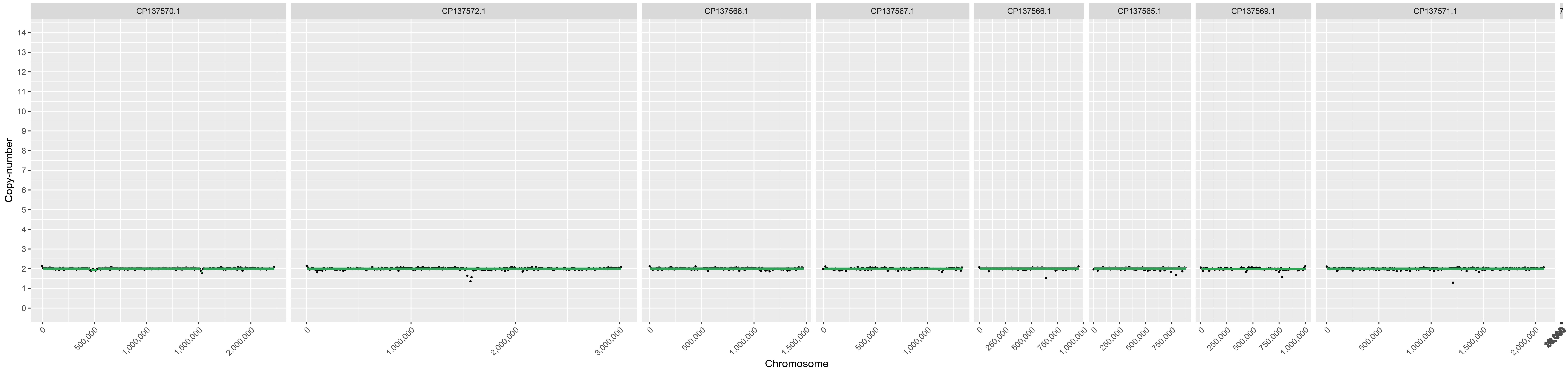

35763_2_42 whole genome

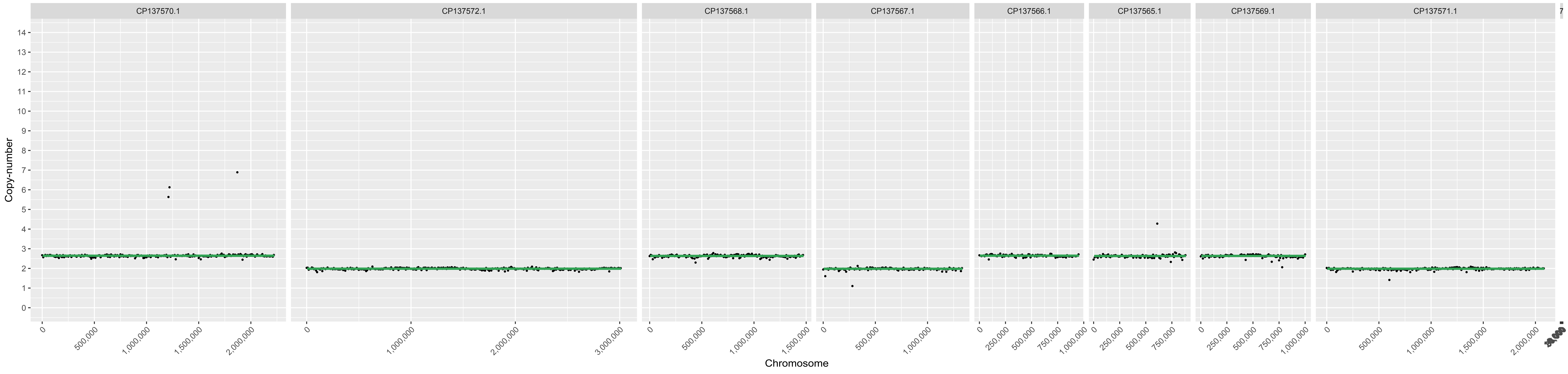

35763_2_43 whole genome

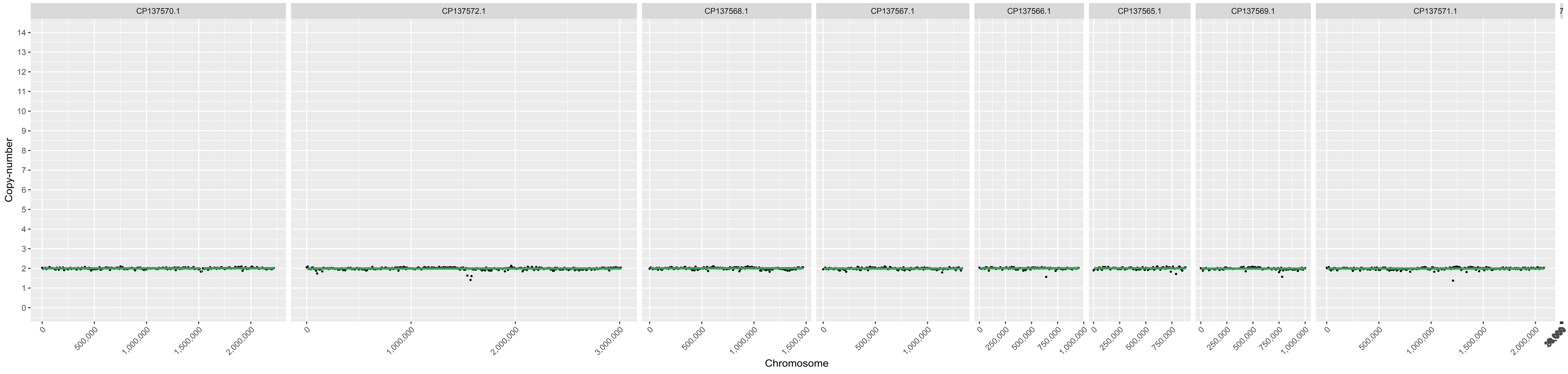

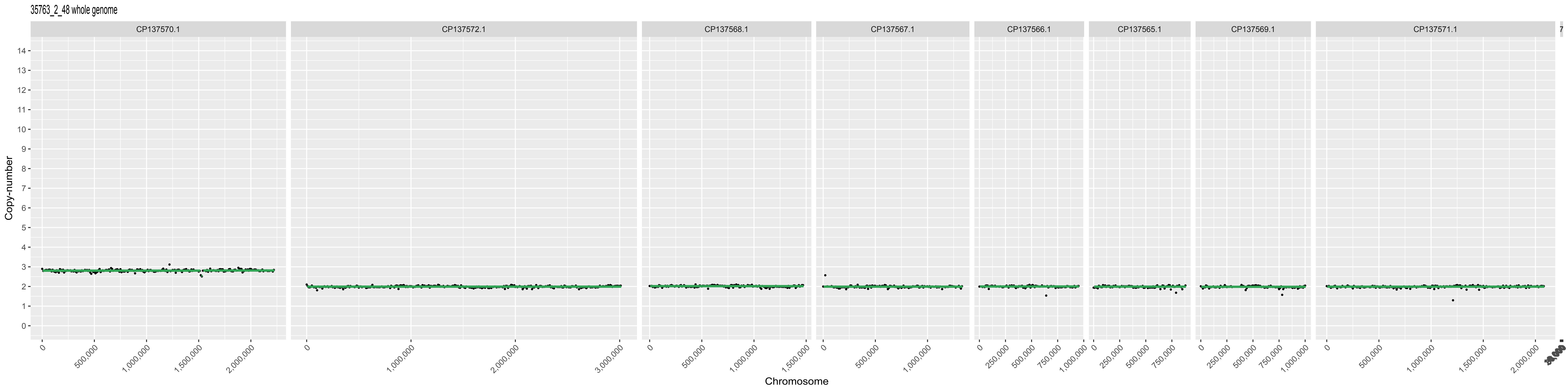

35763_2_48 whole genome
CP137570.1
CP137572.1
CP137568.1
CP137567.1
CP137566.1
CP137565.1
CP137569.1
CP137571.1
Copy-number
Chromosome
500,000
1,000,000
1,500,000
2,000,000
3,000,000
250,000
750,000

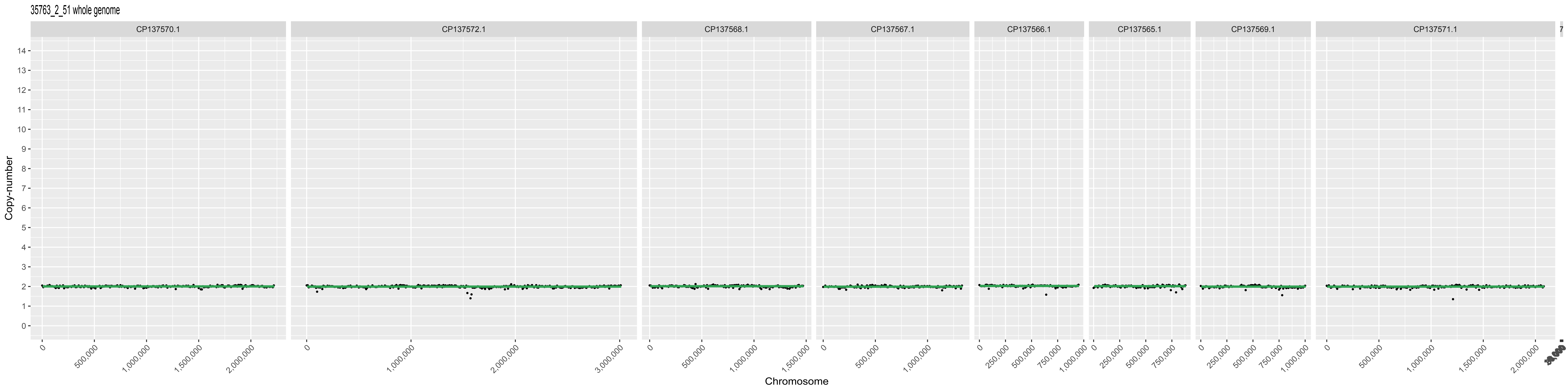
35763_2_51 whole genome
CP137570.1
CP137572.1
CP137568.1
CP137567.1
CP137566.1
CP137565.1
CP137569.1
CP137571.1
Copy-number
Chromosome
500,000
1,000,000
1,500,000
2,000,000
3,000,000
250,000
750,000

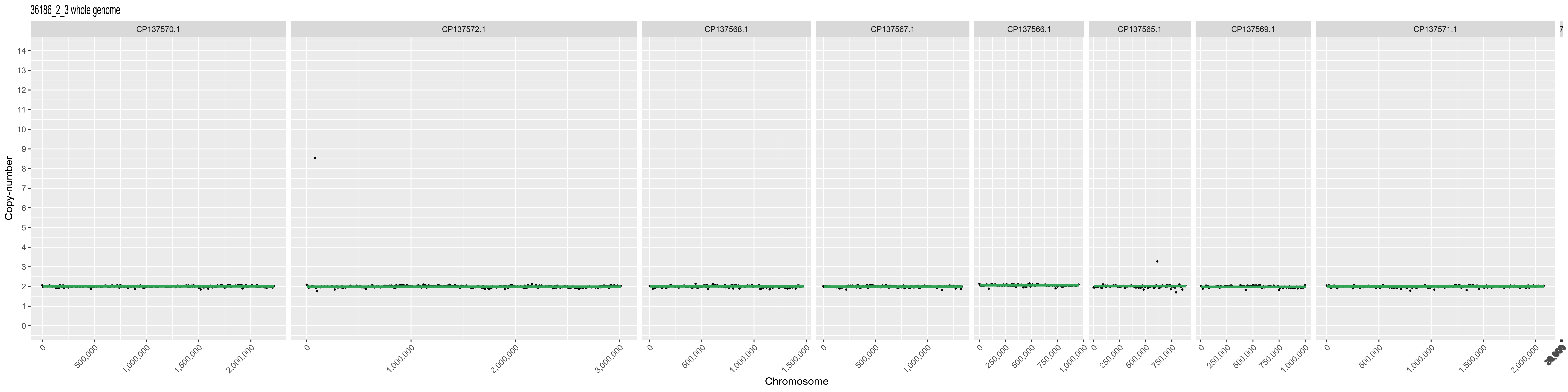

36186_2_3 whole genome
CP137570.1
CP137572.1
CP137568.1
CP137567.1
CP137566.1
CP137565.1
CP137569.1
CP137571.1
Copy-number
Chromosome

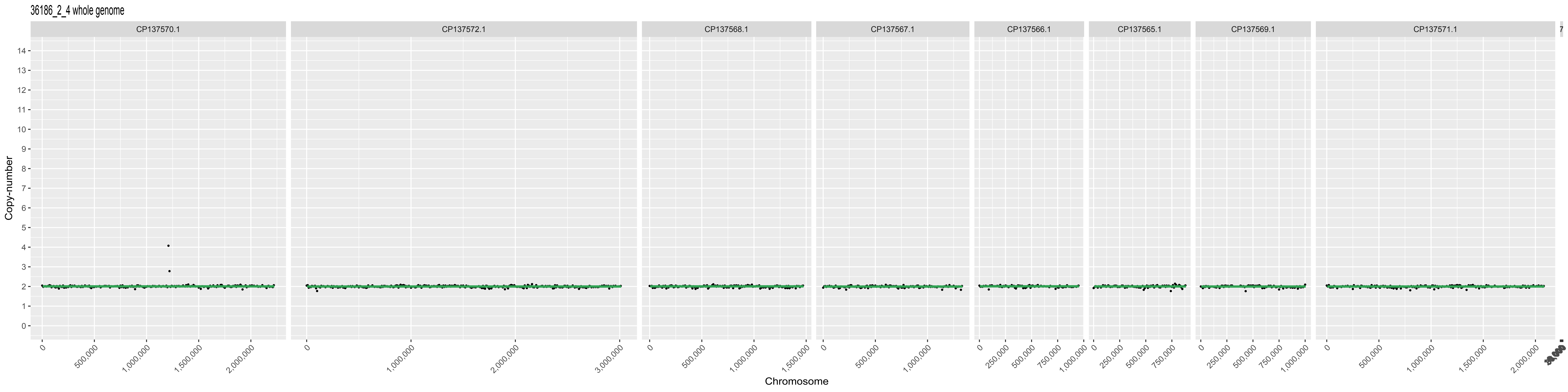
36186_2_4 whole genome
CP137570.1
CP137572.1
CP137568.1
CP137567.1
CP137566.1
CP137565.1
CP137569.1
CP137571.1
Copy-number
Chromosome
500,000
1,000,000
1,500,000
2,000,000
3,000,000
250,000
750,000

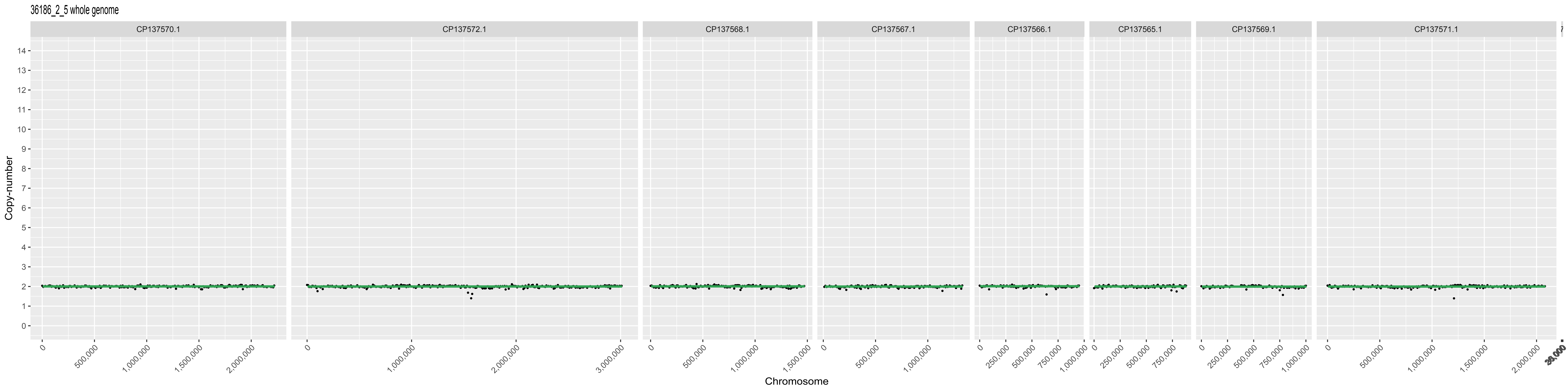
36186_2_5 whole genome
CP137570.1
CP137572.1
CP137568.1
CP137567.1
CP137566.1
CP137565.1
CP137569.1
CP137571.1
Copy-number
Chromosome
0
500,000
1,000,000
1,500,000
2,000,000
3,000,000
250,000
750,000

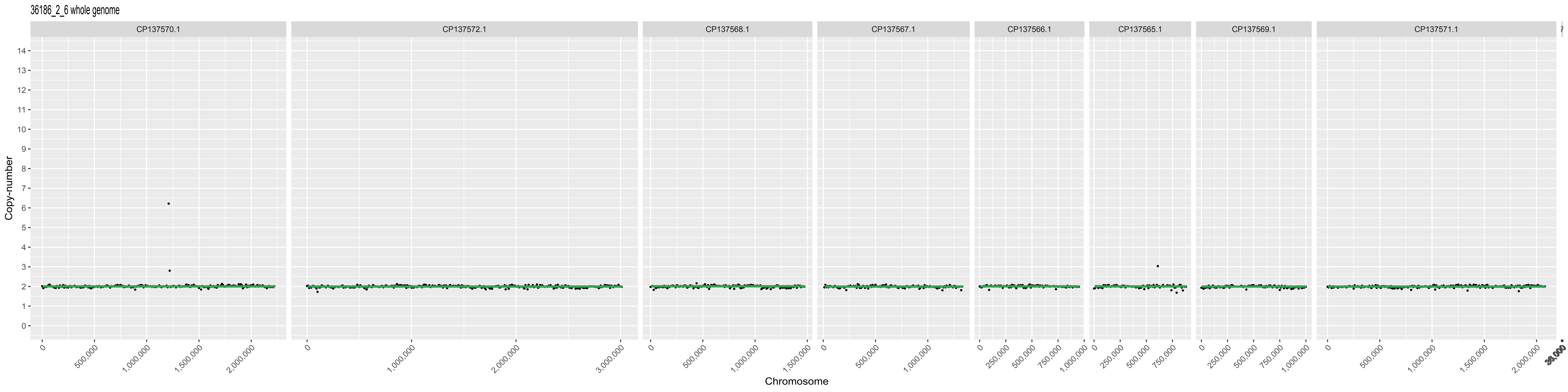
36186_2_6 whole genome
CP137570.1
CP137572.1
CP137568.1
CP137567.1
CP137566.1
CP137565.1
CP137569.1
CP137571.1
Copy-number
Chromosome

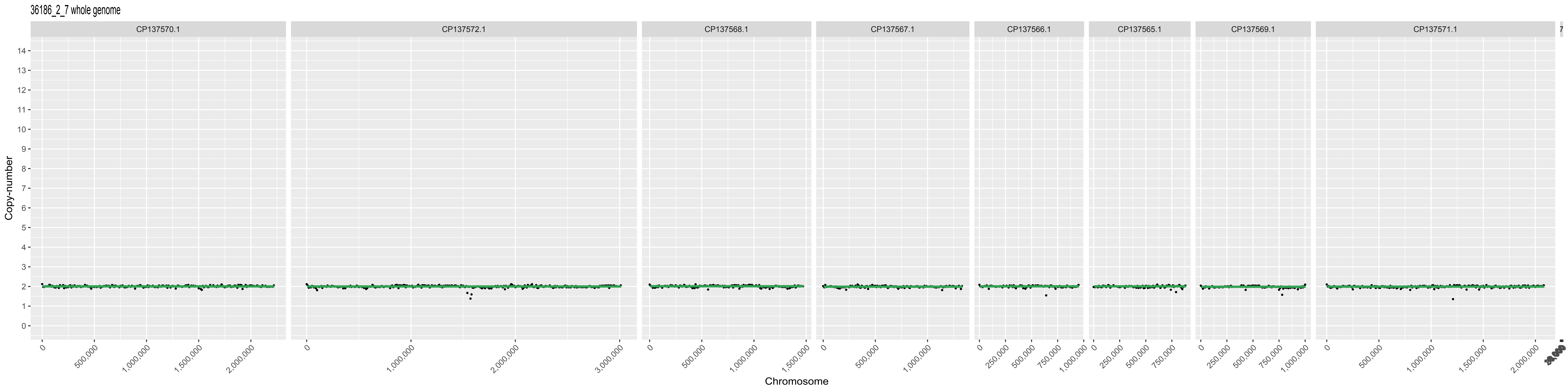
36186_2_7 whole genome
CP137570.1
CP137572.1
CP137568.1
CP137567.1
CP137566.1
CP137565.1
CP137569.1
CP137571.1
Copy-number
Chromosome

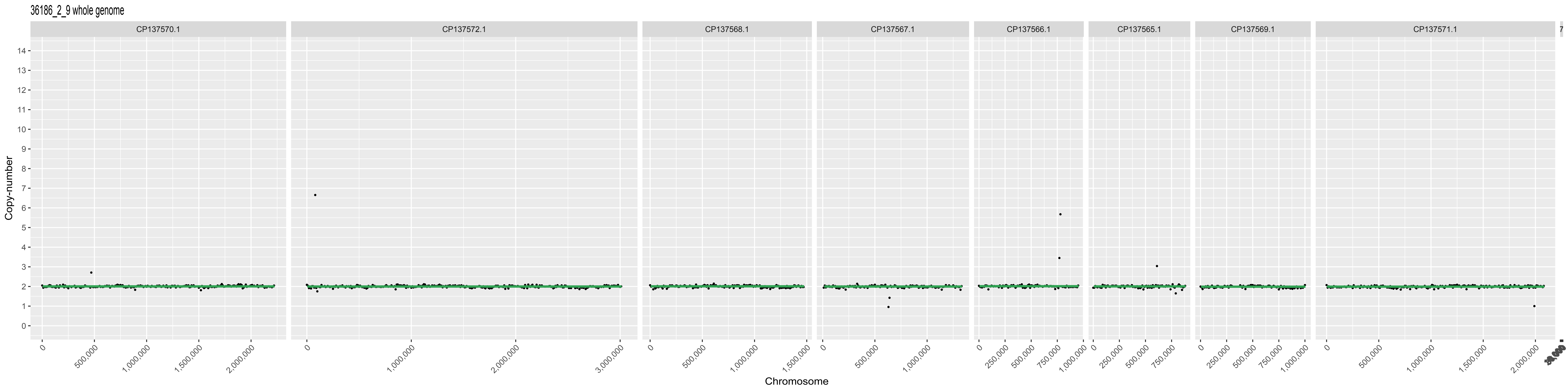
36186_2_9 whole genome
CP137570.1
CP137572.1
CP137568.1
CP137567.1
CP137566.1
CP137565.1
CP137569.1
CP137571.1
Copy-number
0
500,000
1,000,000
1,500,000
2,000,000
3,000,000
250,000
750,000
Chromosome

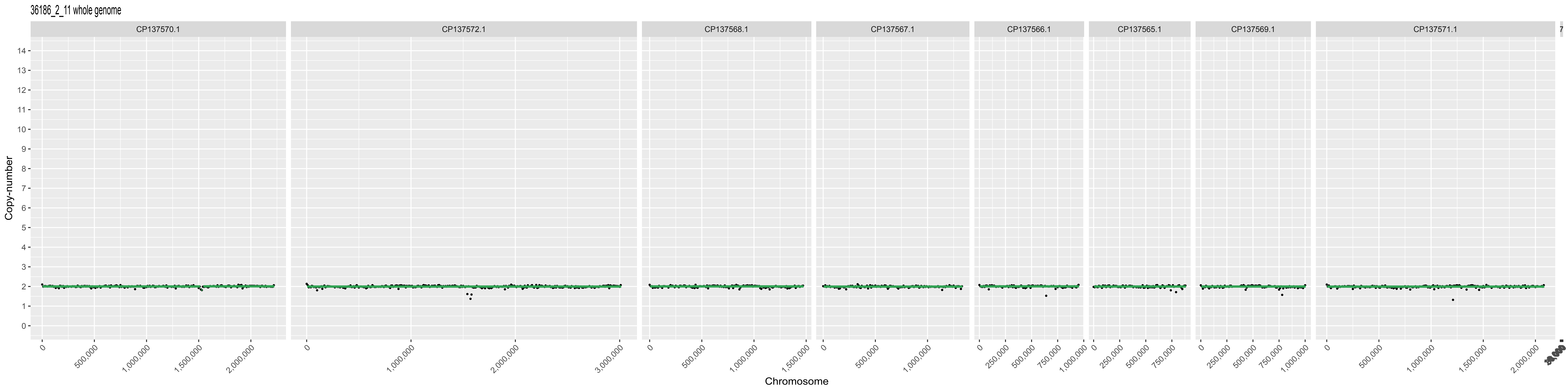
36186_2_11 whole genome
CP137570.1
CP137572.1
CP137568.1
CP137567.1
CP137566.1
CP137565.1
CP137569.1
CP137571.1
Copy-number
Chromosome
500,000
1,000,000
1,500,000
2,000,000
3,000,000
250,000
750,000

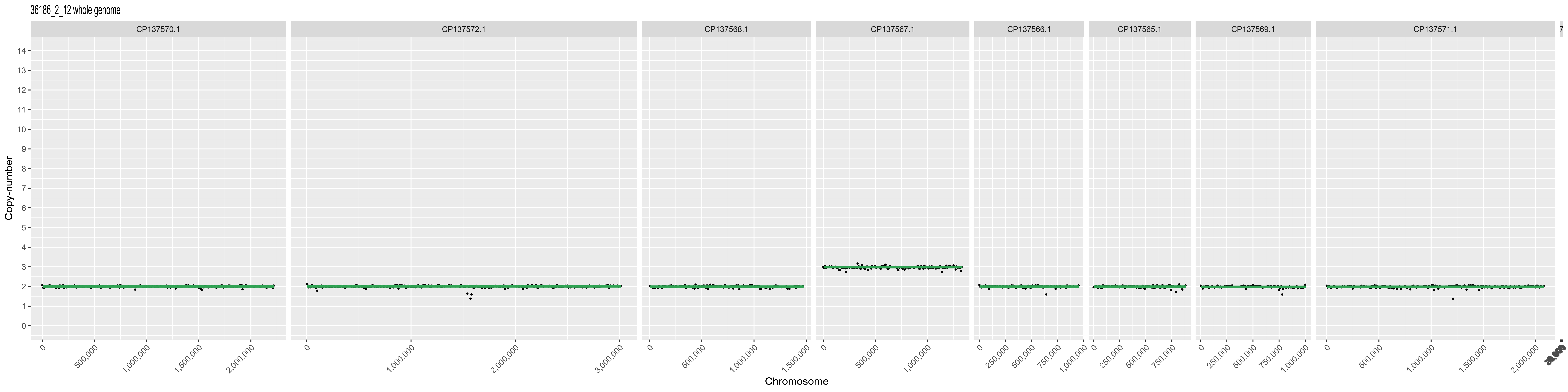

36186_2_12 whole genome
CP137570.1
CP137572.1
CP137568.1
CP137567.1
CP137566.1
CP137565.1
CP137569.1
CP137571.1
Copy-number
Chromosome

36186_2_14 whole genome

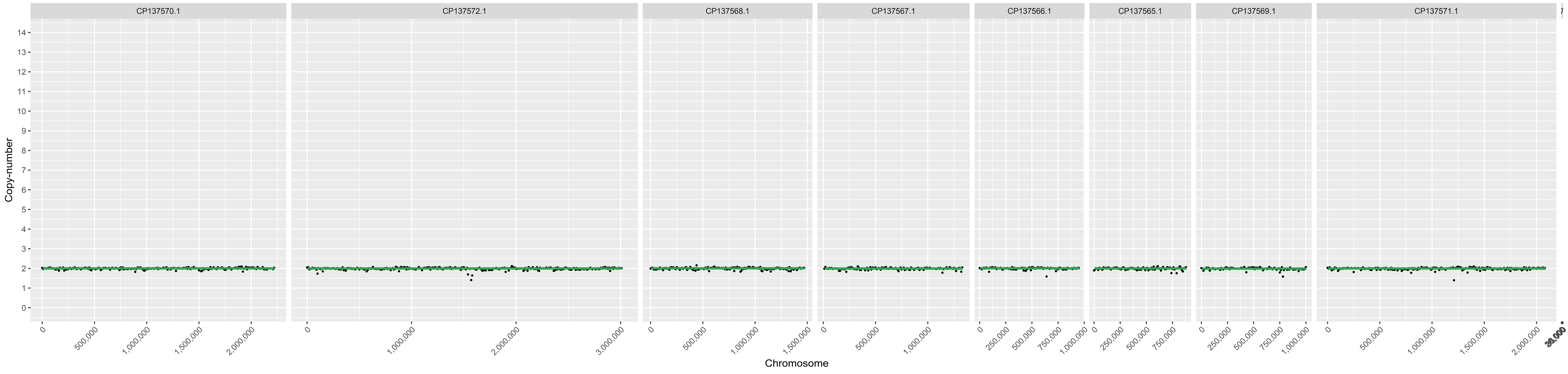

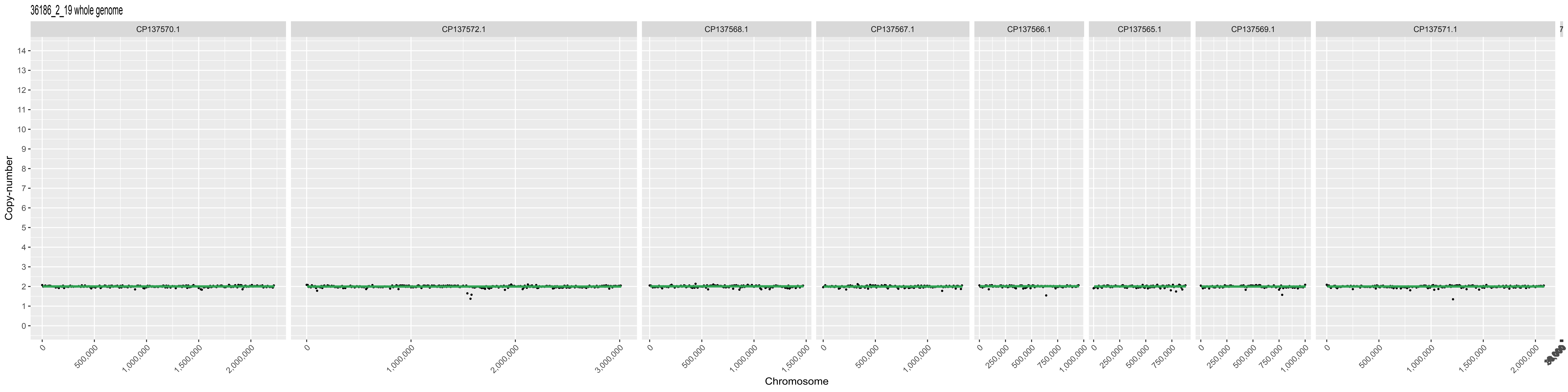

36186_2_19 whole genome
CP137570.1
CP137572.1
CP137568.1
CP137567.1
CP137566.1
CP137565.1
CP137569.1
CP137571.1
Copy-number
Chromosome

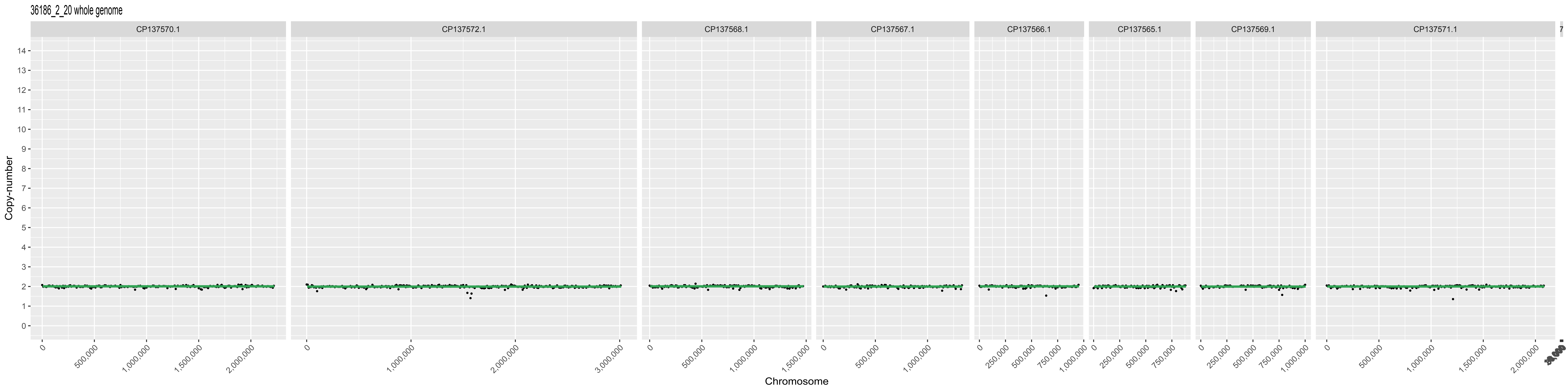
36186_2_20 whole genome
CP137570.1
CP137572.1
CP137568.1
CP137567.1
CP137566.1
CP137565.1
CP137569.1
CP137571.1
Copy-number
Chromosome

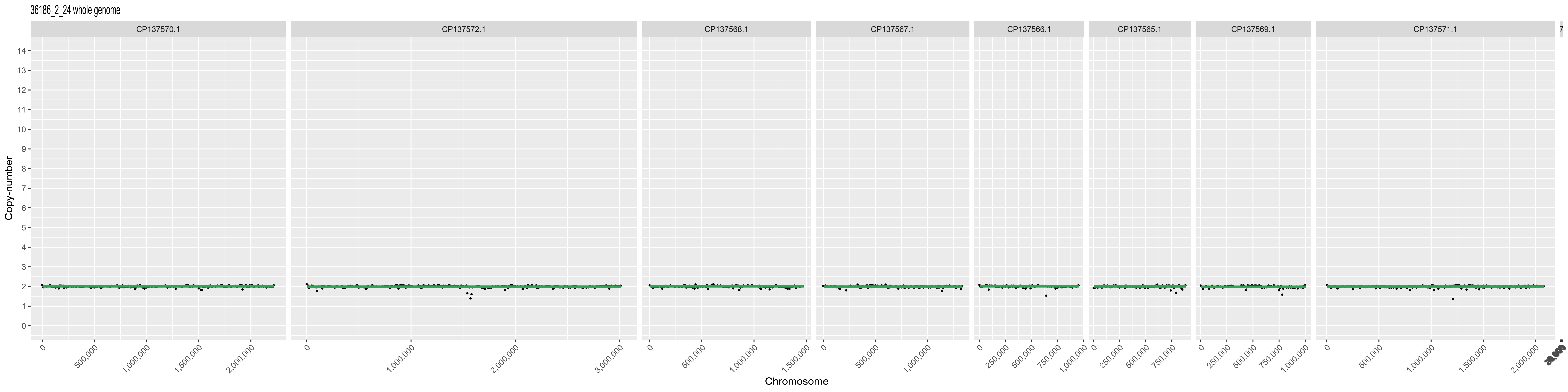

36186_2_24 whole genome
CP137570.1
CP137572.1
CP137568.1
CP137567.1
CP137566.1
CP137565.1
CP137569.1
CP137571.1
Copy-number
Chromosome

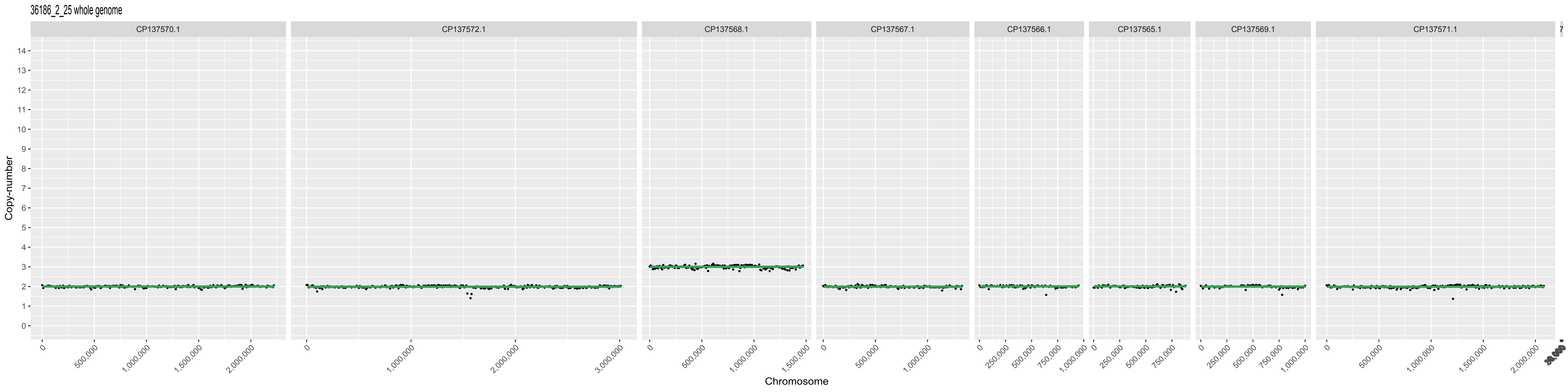
36186_2_25 whole genome
CP137570.1
CP137572.1
CP137568.1
CP137567.1
CP137566.1
CP137565.1
CP137569.1
CP137571.1
Copy-number
Chromosome

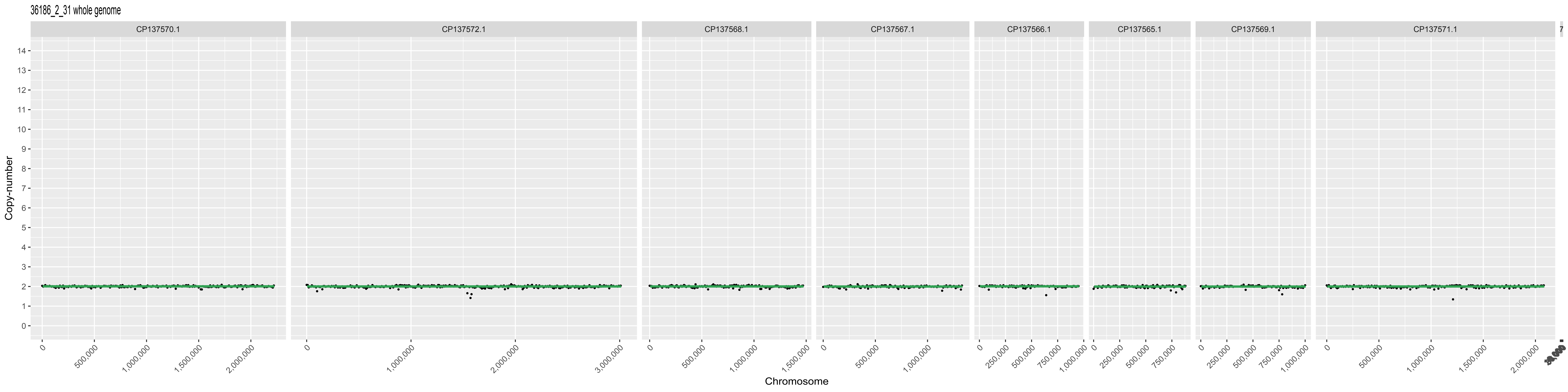
36186_2_31 whole genome
CP137570.1
CP137572.1
CP137568.1
CP137567.1
CP137566.1
CP137565.1
CP137569.1
CP137571.1
Copy-number
Chromosome

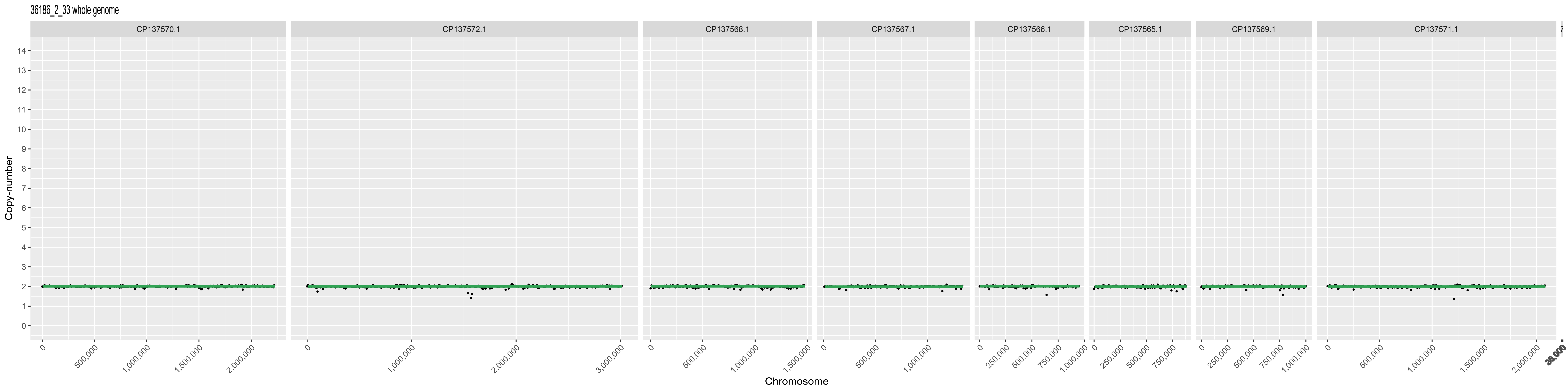

36186_2_33 whole genome
CP137570.1
CP137572.1
CP137568.1
CP137567.1
CP137566.1
CP137565.1
CP137569.1
CP137571.1
Copy-number
Chromosome

36186_2_35 whole genome

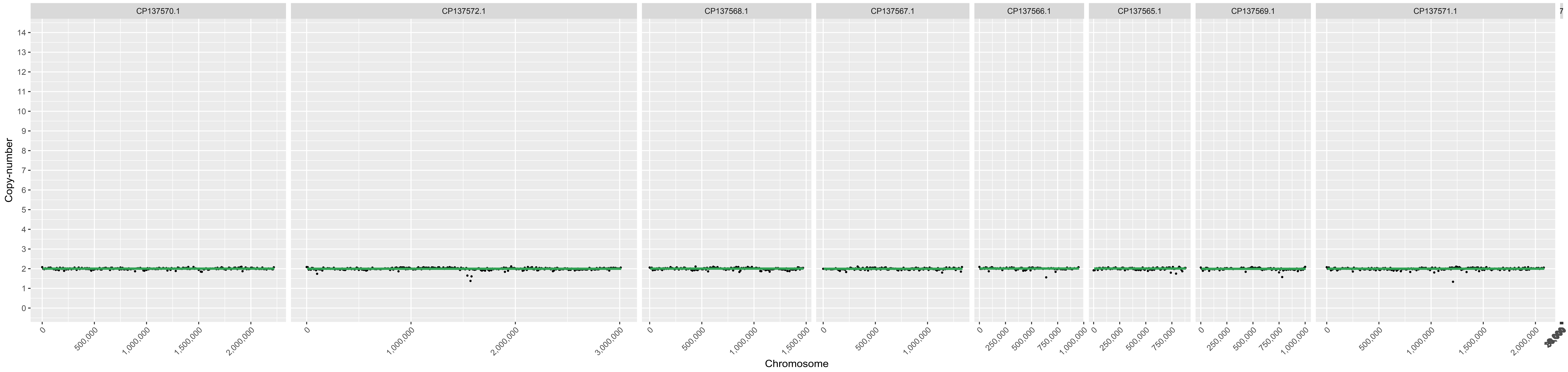

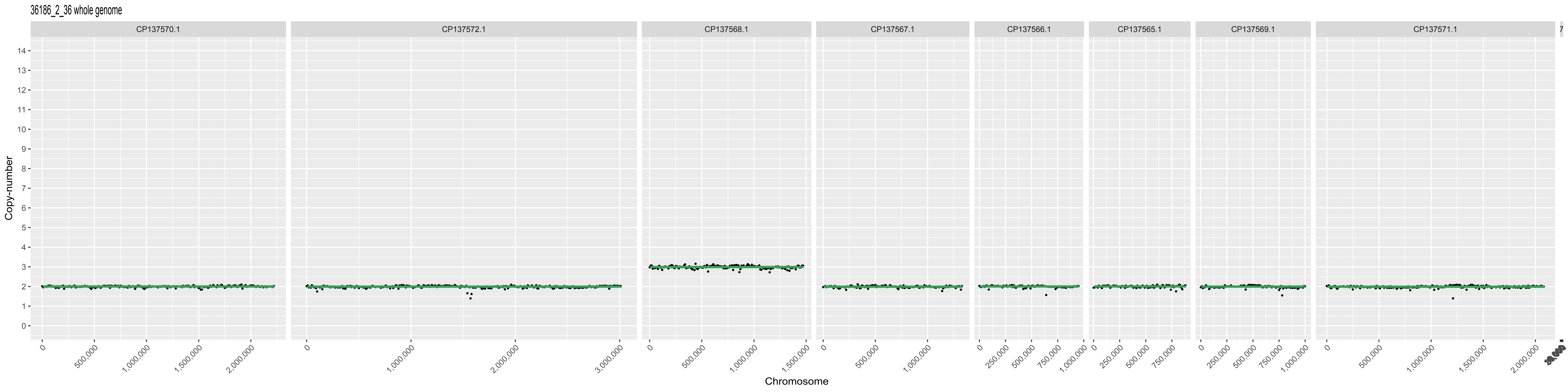
36186_2_36 whole genome
CP137570.1
CP137572.1
CP137568.1
CP137567.1
CP137566.1
CP137565.1
CP137569.1
CP137571.1
Copy-number
Chromosome
500,000
1,000,000
1,500,000
2,000,000
3,000,000
250,000
750,000

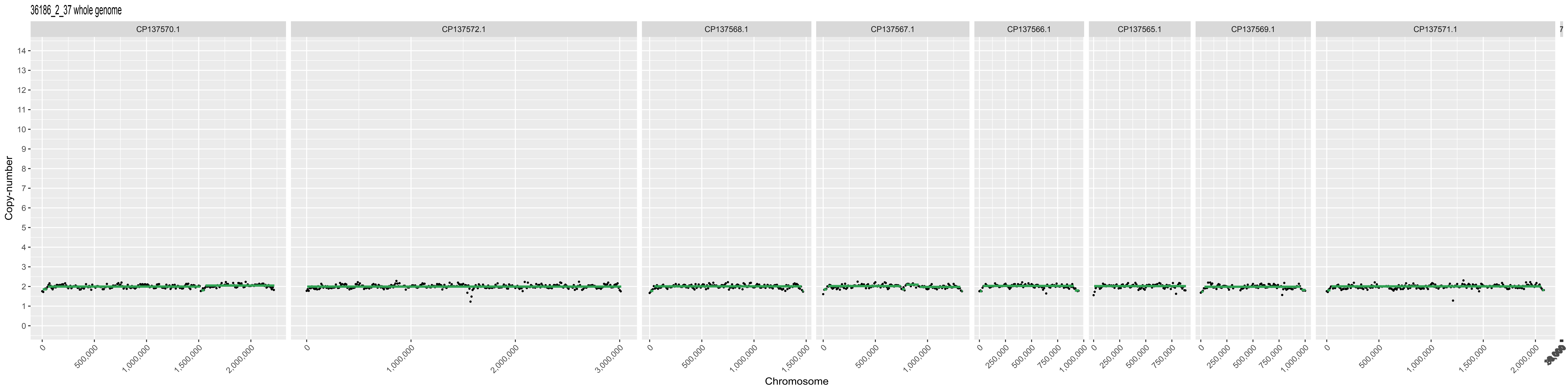
36186_2_37 whole genome
CP137570.1
CP137572.1
CP137568.1
CP137567.1
CP137566.1
CP137565.1
CP137569.1
CP137571.1
Copy-number
Chromosome
0
500,000
1,000,000
1,500,000
2,000,000
3,000,000
250,000
750,000

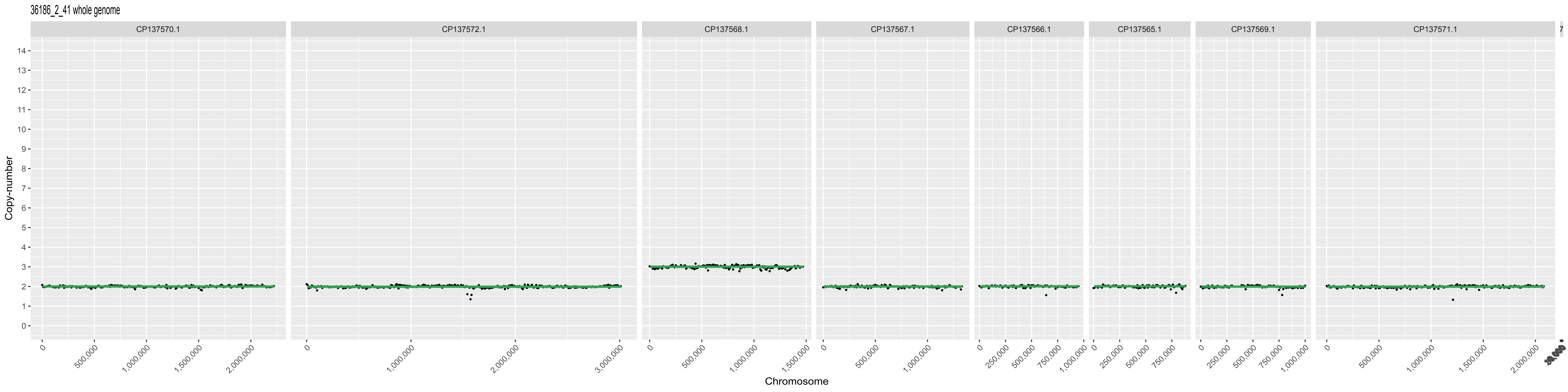
36186_2_41 whole genome
CP137570.1
CP137572.1
CP137568.1
CP137567.1
CP137566.1
CP137565.1
CP137569.1
CP137571.1
Copy-number
Chromosome

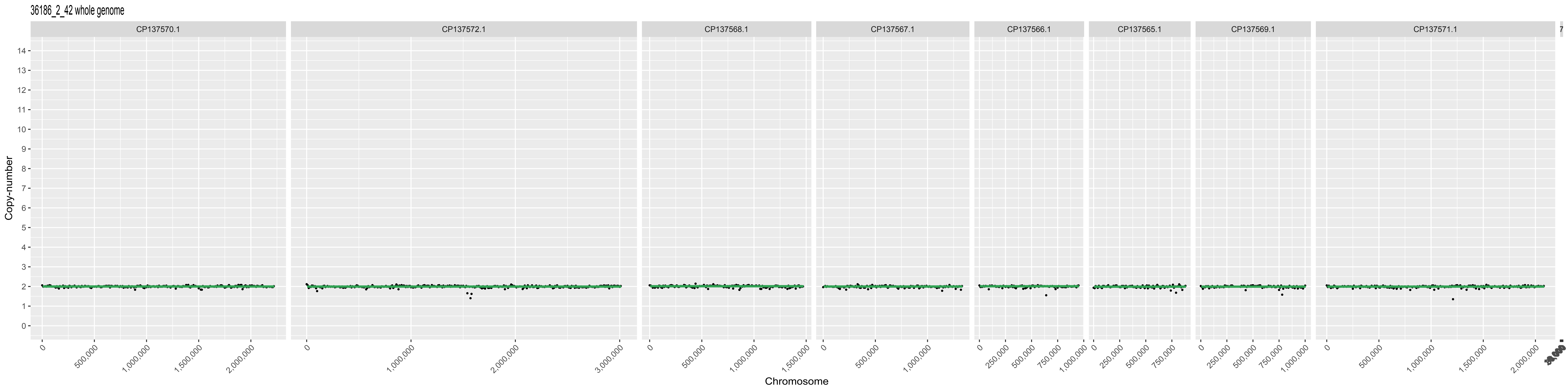

36186_2_42 whole genome
CP137570.1
CP137572.1
CP137568.1
CP137567.1
CP137566.1
CP137565.1
CP137569.1
CP137571.1
Copy-number
Chromosome

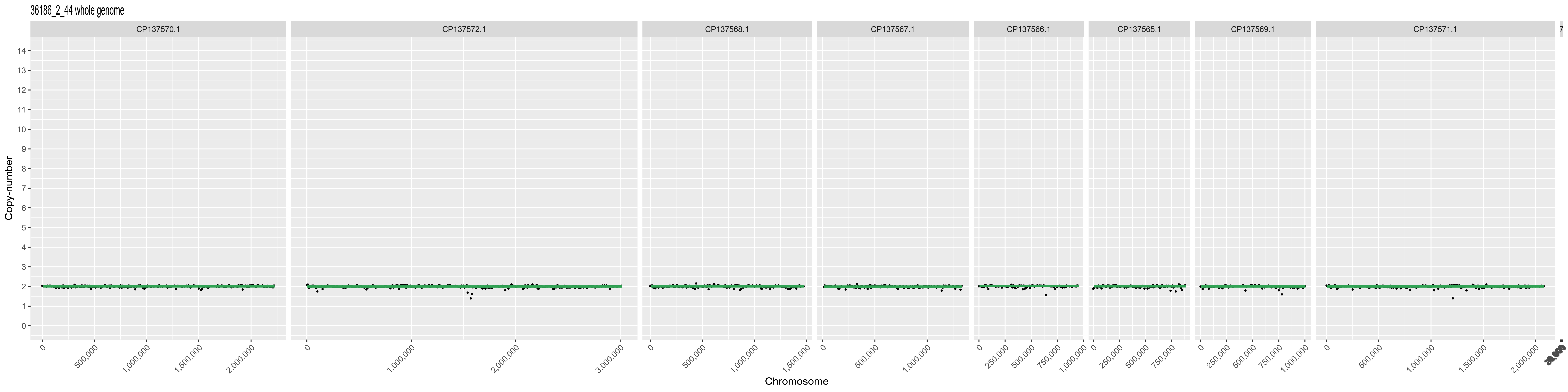
36186_2_44 whole genome
CP137570.1
CP137572.1
CP137568.1
CP137567.1
CP137566.1
CP137565.1
CP137569.1
CP137571.1
Copy-number
Chromosome

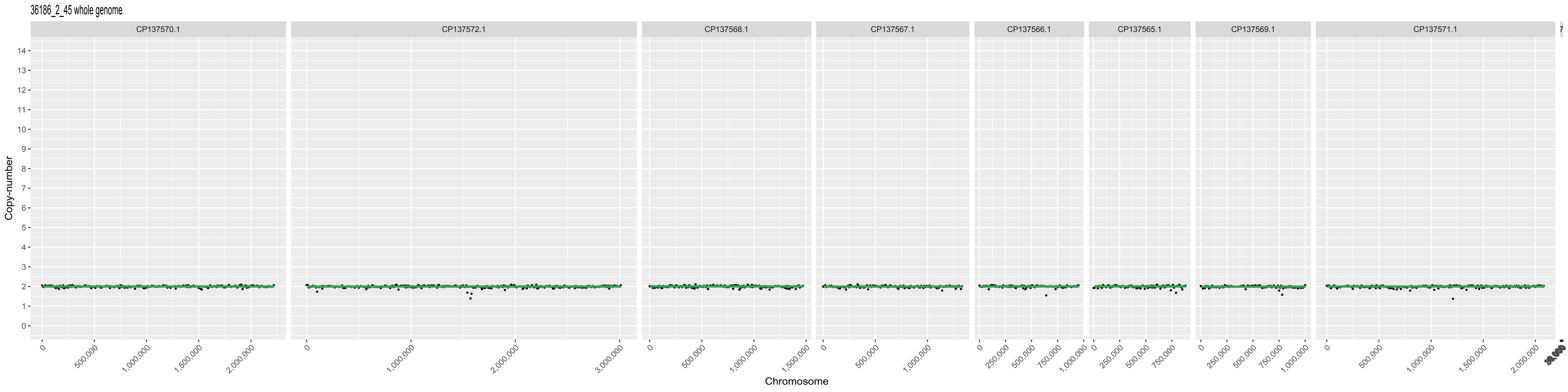

36186_2_45 whole genome
CP137570.1
CP137572.1
CP137568.1
CP137567.1
CP137566.1
CP137565.1
CP137569.1
CP137571.1
Copy-number
Chromosome

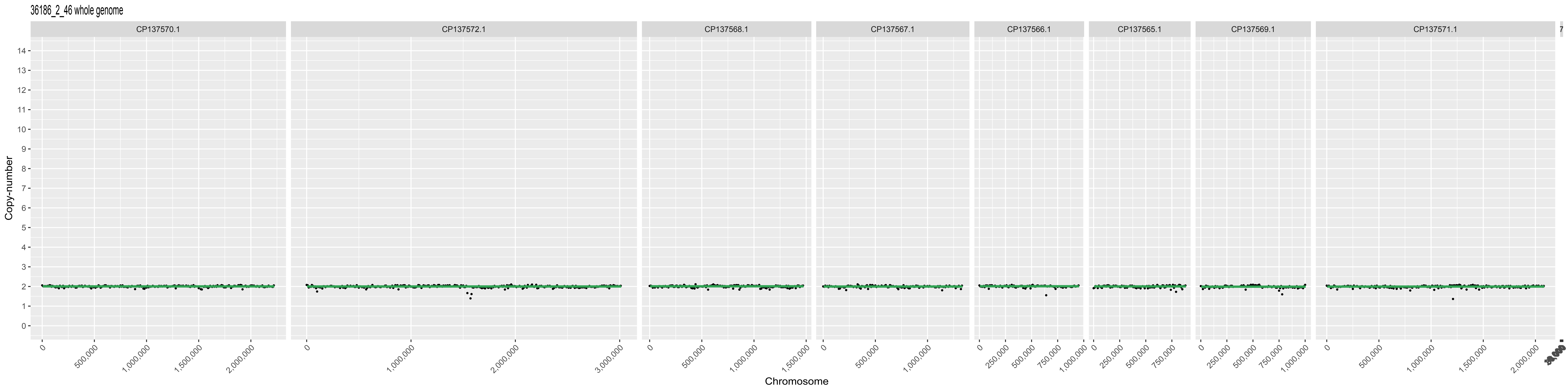
36186_2_46 whole genome
CP137570.1
CP137572.1
CP137568.1
CP137567.1
CP137566.1
CP137565.1
CP137569.1
CP137571.1
Copy-number
Chromosome

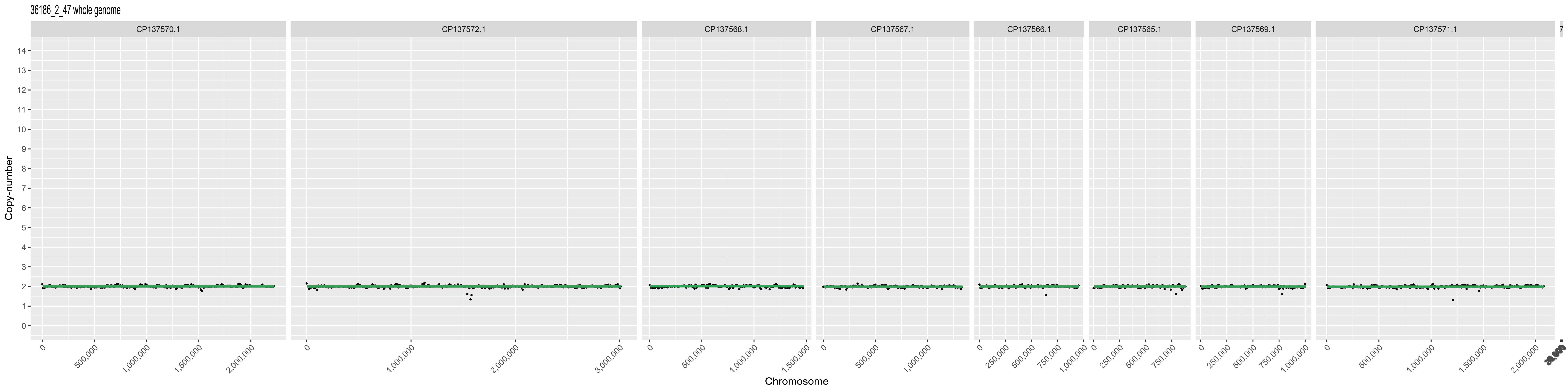

36186_2_47 whole genome
CP137570.1
CP137572.1
CP137568.1
CP137567.1
CP137566.1
CP137565.1
CP137569.1
CP137571.1
Copy-number
Chromosome

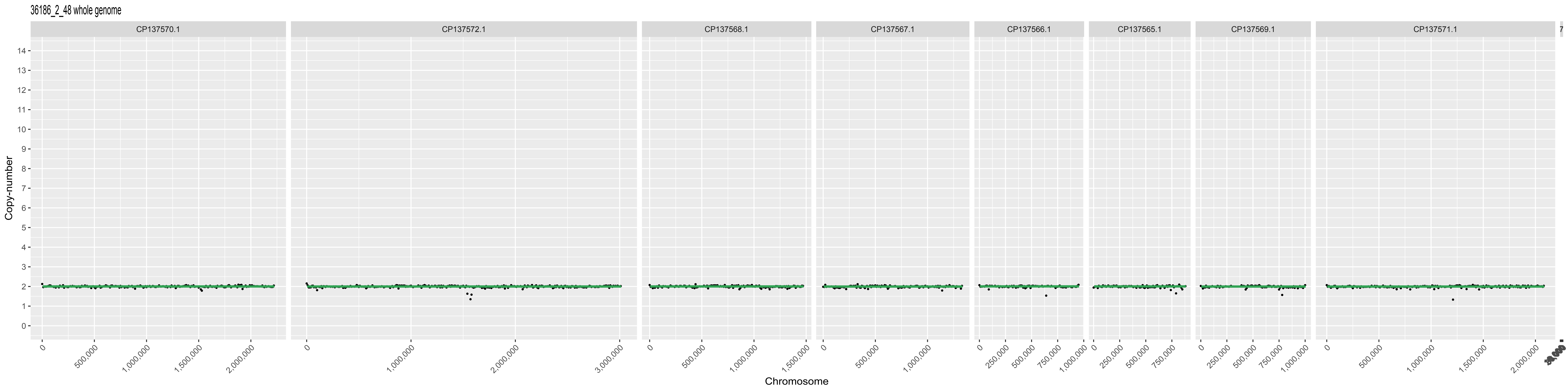

36186_2_48 whole genome
CP137570.1
CP137572.1
CP137568.1
CP137567.1
CP137566.1
CP137565.1
CP137569.1
CP137571.1
Copy-number
Chromosome

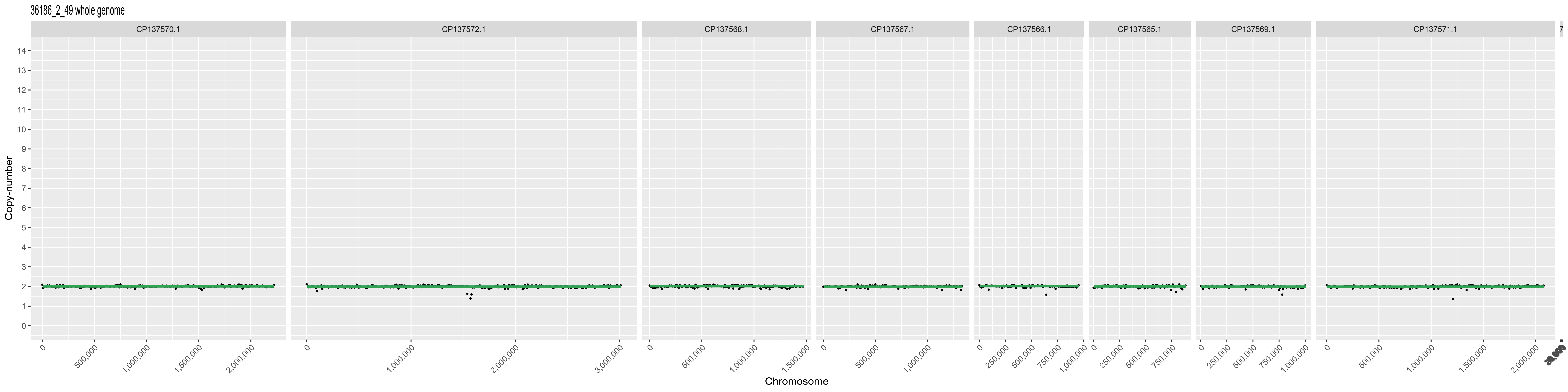

36186_2_49 whole genome
CP137570.1
CP137572.1
CP137568.1
CP137567.1
CP137566.1
CP137565.1
CP137569.1
CP137571.1
Copy-number
Chromosome

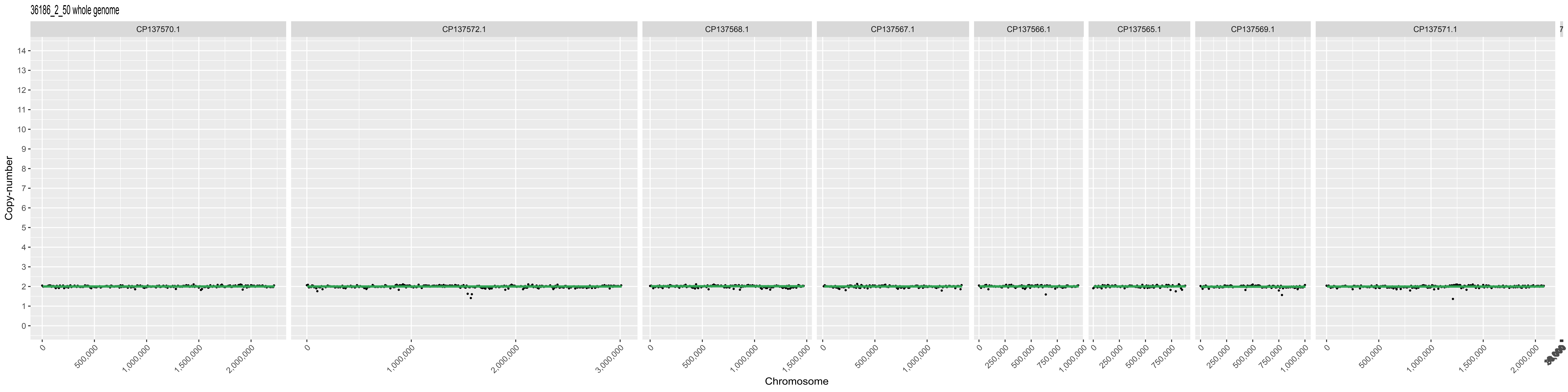
36186_2_50 whole genome
CP137570.1
CP137572.1
CP137568.1
CP137567.1
CP137566.1
CP137565.1
CP137569.1
CP137571.1
Copy-number
Chromosome
500,000
1,000,000
1,500,000
2,000,000
3,000,000
250,000
750,000

36186_2_51 whole genome

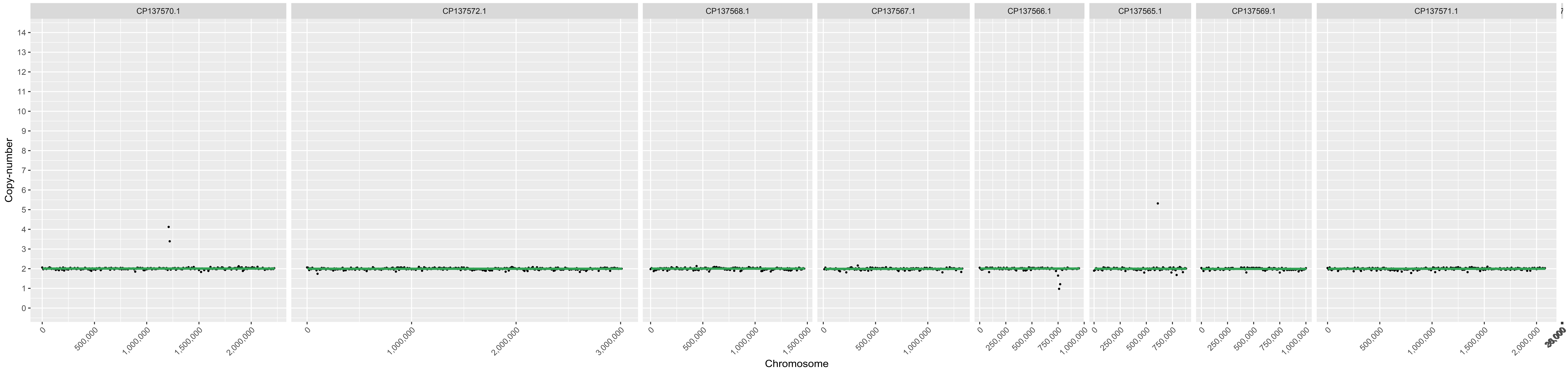

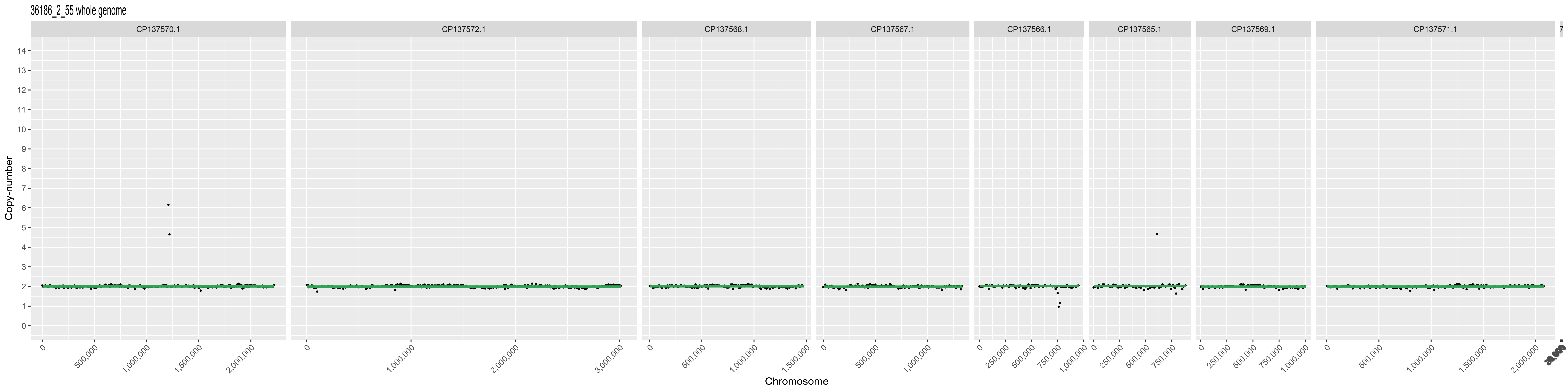
36186_2_55 whole genome
CP137570.1
CP137572.1
CP137568.1
CP137567.1
CP137566.1
CP137565.1
CP137569.1
CP137571.1
Copy-number
Chromosome

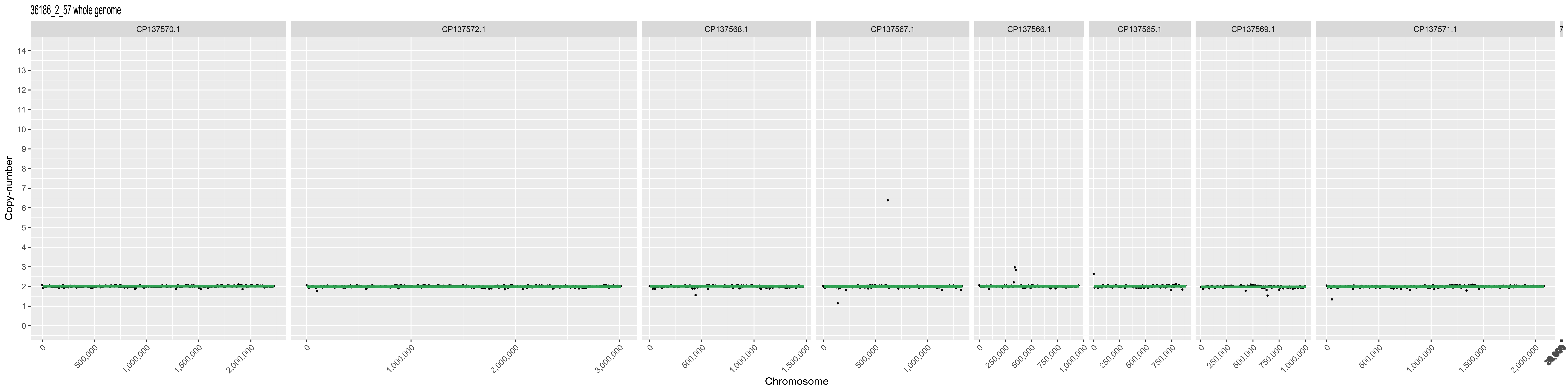

36186_2_57 whole genome
CP137570.1
CP137572.1
CP137568.1
CP137567.1
CP137566.1
CP137565.1
CP137569.1
CP137571.1
Copy-number
Chromosome

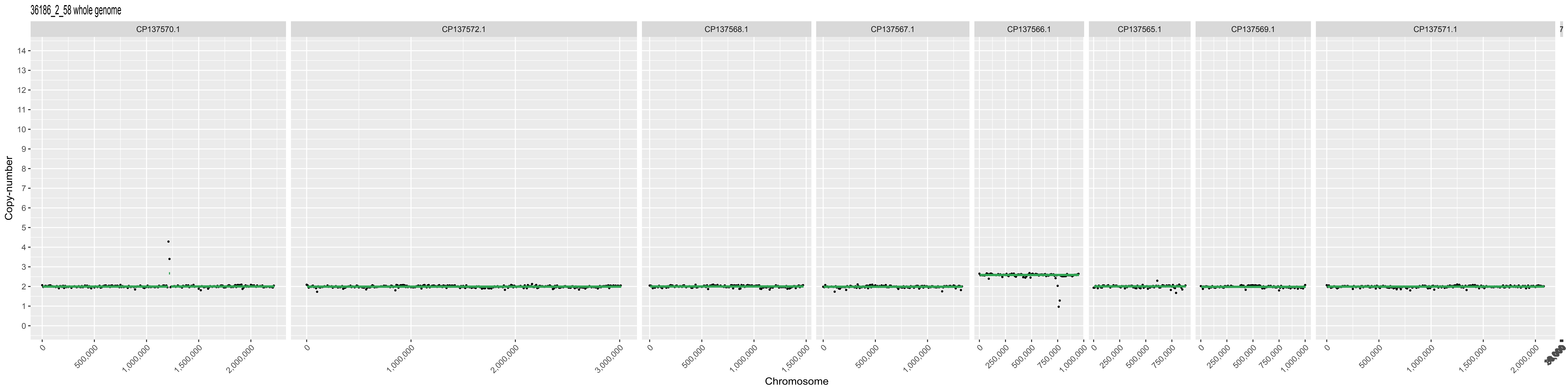

36186_2_58 whole genome
CP137570.1
CP137572.1
CP137568.1
CP137567.1
CP137566.1
CP137565.1
CP137569.1
CP137571.1
Copy-number
Chromosome

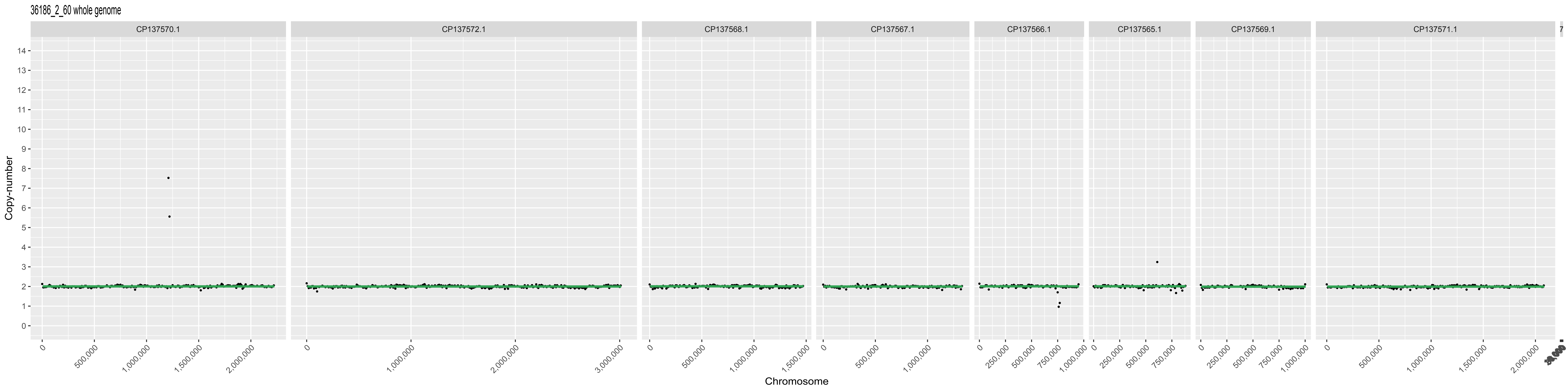

36186_2_60 whole genome
CP137570.1
CP137572.1
CP137568.1
CP137567.1
CP137566.1
CP137565.1
CP137569.1
CP137571.1
Copy-number
Chromosome

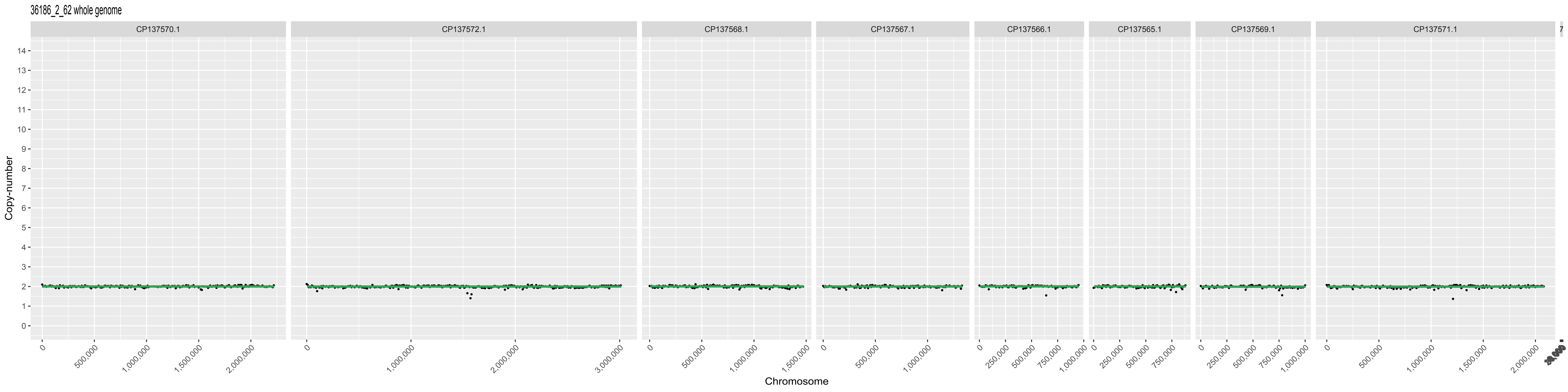
36186_2_62 whole genome
CP137570.1
CP137572.1
CP137568.1
CP137567.1
CP137566.1
CP137565.1
CP137569.1
CP137571.1
Copy-number
Chromosome

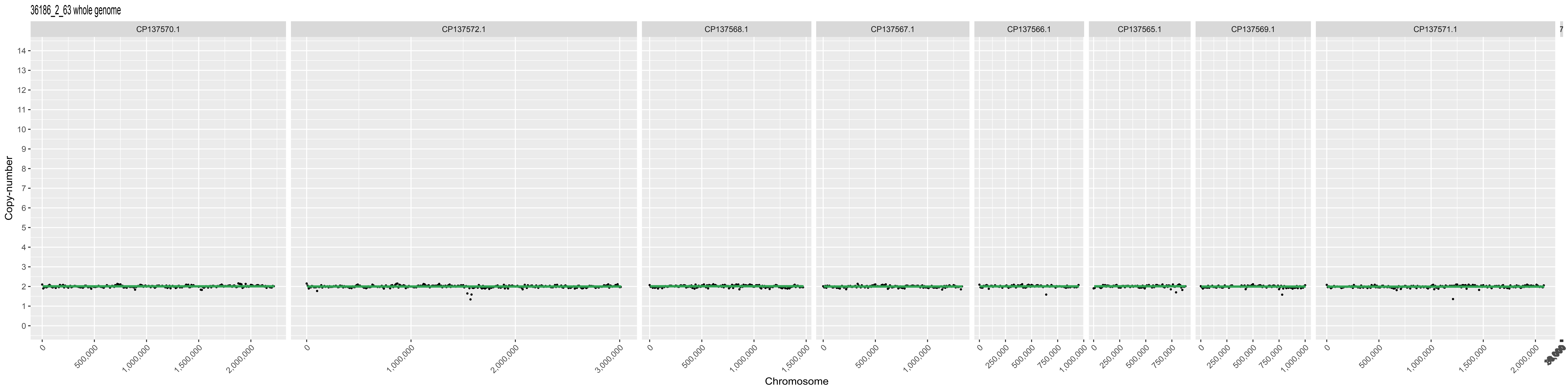

36186_2_63 whole genome
CP137570.1
CP137572.1
CP137568.1
CP137567.1
CP137566.1
CP137565.1
CP137569.1
CP137571.1
Copy-number
Chromosome
0
500,000
1,000,000
1,500,000
2,000,000
3,000,000
250,000
750,000

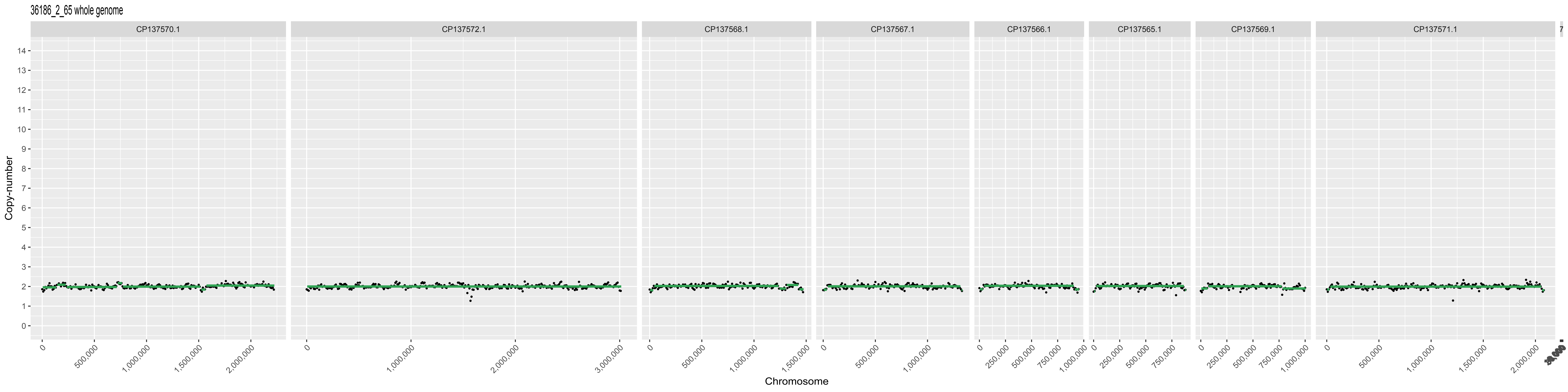

36186_2_65 whole genome
CP137570.1
CP137572.1
CP137568.1
CP137567.1
CP137566.1
CP137565.1
CP137569.1
CP137571.1
Copy-number
Chromosome

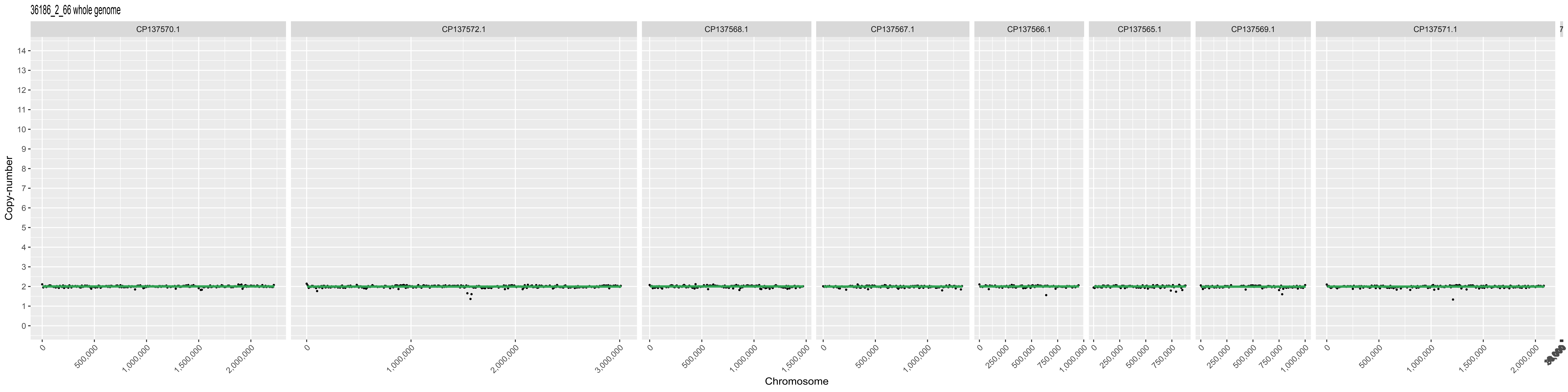

36186_2_66 whole genome
CP137570.1
CP137572.1
CP137568.1
CP137567.1
CP137566.1
CP137565.1
CP137569.1
CP137571.1
Copy-number
Chromosome
500,000
1,000,000
1,500,000
2,000,000
3,000,000
250,000
750,000

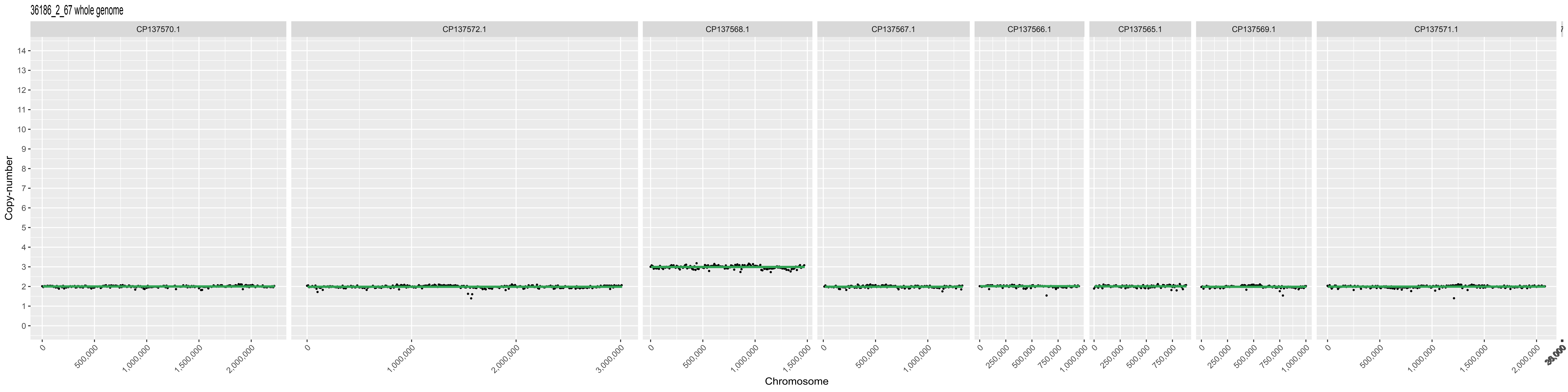

36186_2_67 whole genome
CP137570.1
CP137572.1
CP137568.1
CP137567.1
CP137566.1
CP137565.1
CP137569.1
CP137571.1
Copy-number
Chromosome

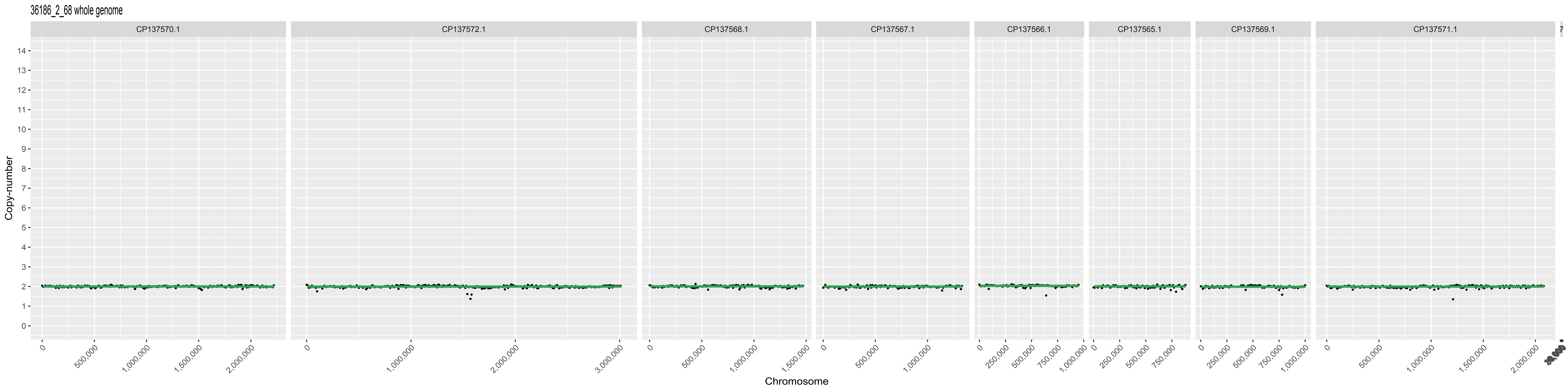

36186_2_68 whole genome
CP137570.1
CP137572.1
CP137568.1
CP137567.1
CP137566.1
CP137565.1
CP137569.1
CP137571.1
Copy-number
Chromosome

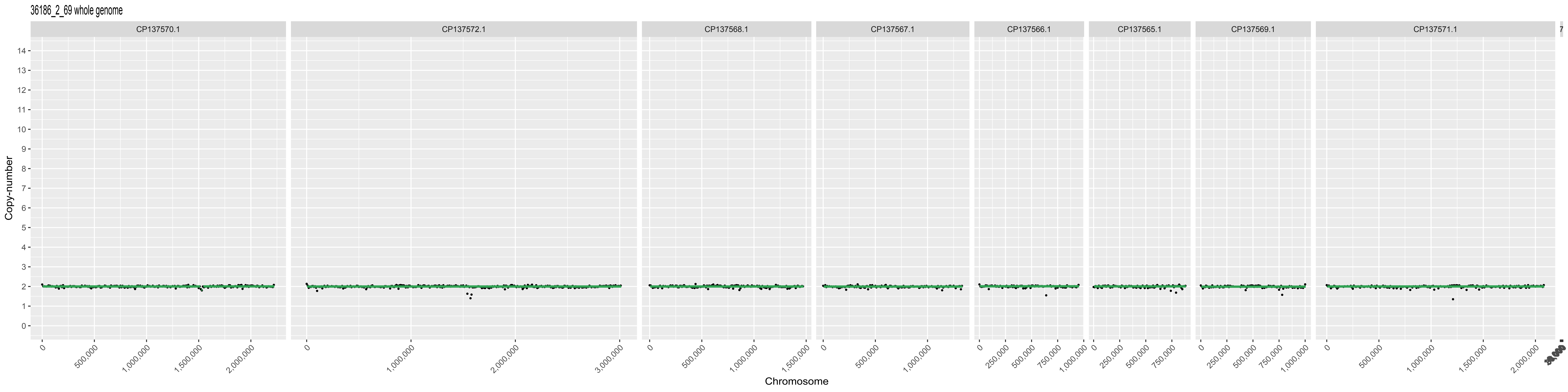

36186_2_69 whole genome
CP137570.1
CP137572.1
CP137568.1
CP137567.1
CP137566.1
CP137565.1
CP137569.1
CP137571.1
Copy-number
Chromosome

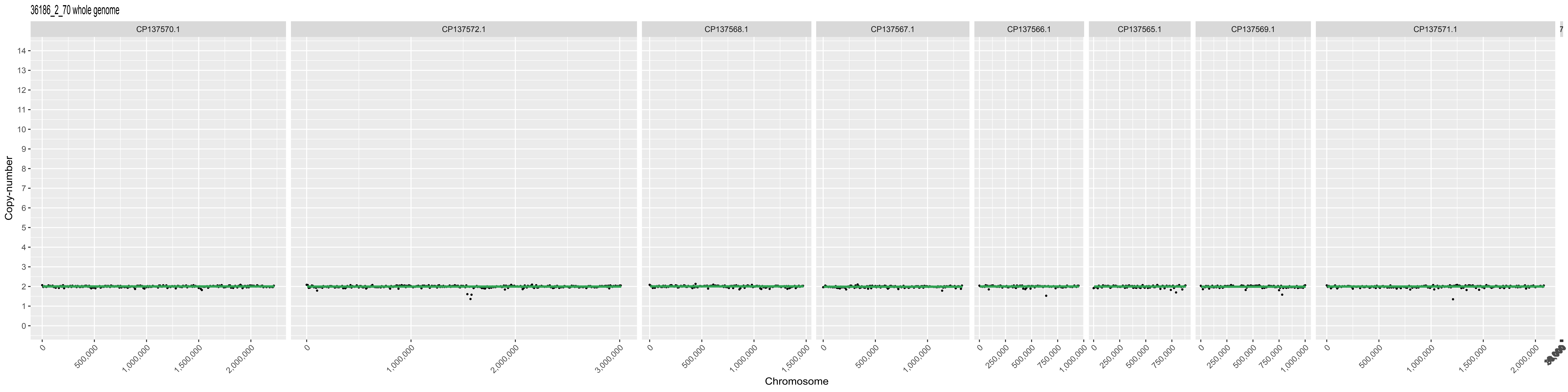

36186_2_70 whole genome
CP137570.1
CP137572.1
CP137568.1
CP137567.1
CP137566.1
CP137565.1
CP137569.1
CP137571.1
Copy-number
Chromosome

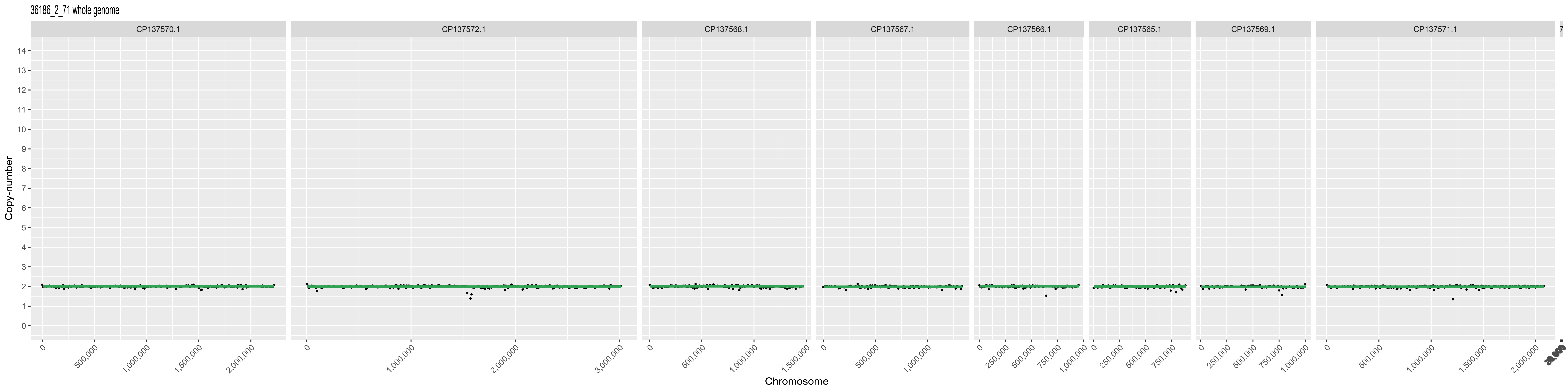
36186_2_71 whole genome
Copy-number
Chromosome
CP137570.1
CP137572.1
CP137568.1
CP137567.1
CP137566.1
CP137565.1
CP137569.1
CP137571.1
0
500,000
1,000,000
1,500,000
2,000,000
3,000,000
250,000
750,000

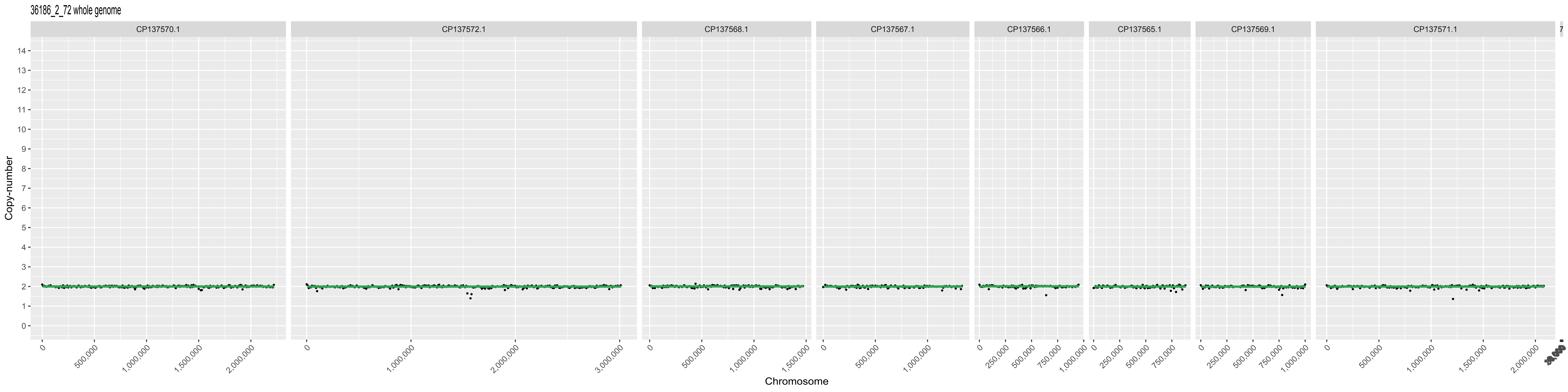
36186_2_72 whole genome
CP137570.1
CP137572.1
CP137568.1
CP137567.1
CP137566.1
CP137565.1
CP137569.1
CP137571.1
Copy-number
Chromosome
500,000
1,000,000
1,500,000
2,000,000
3,000,000
250,000
750,000

36186_2_73 whole genome

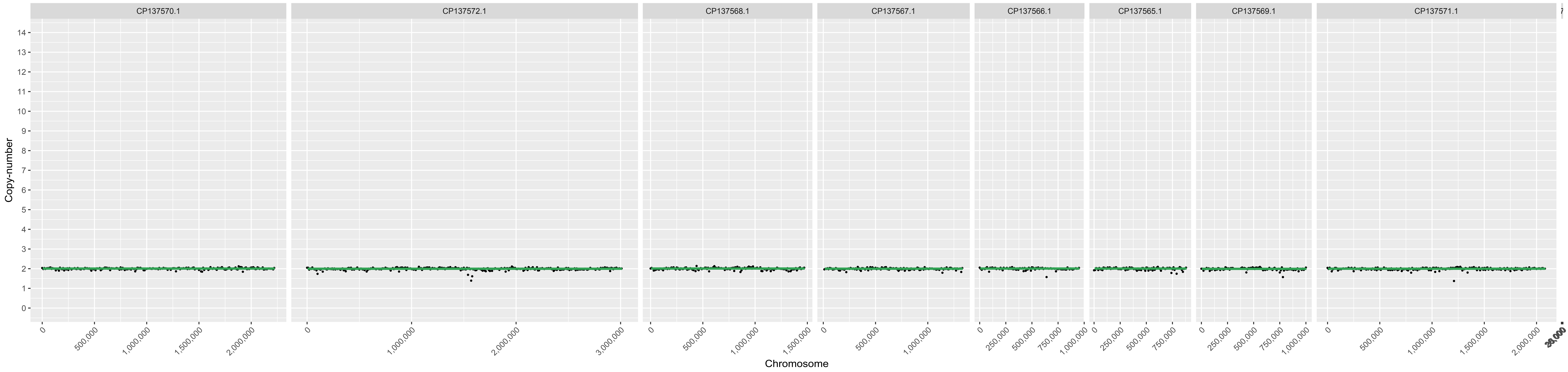

36186_2_74 whole genome

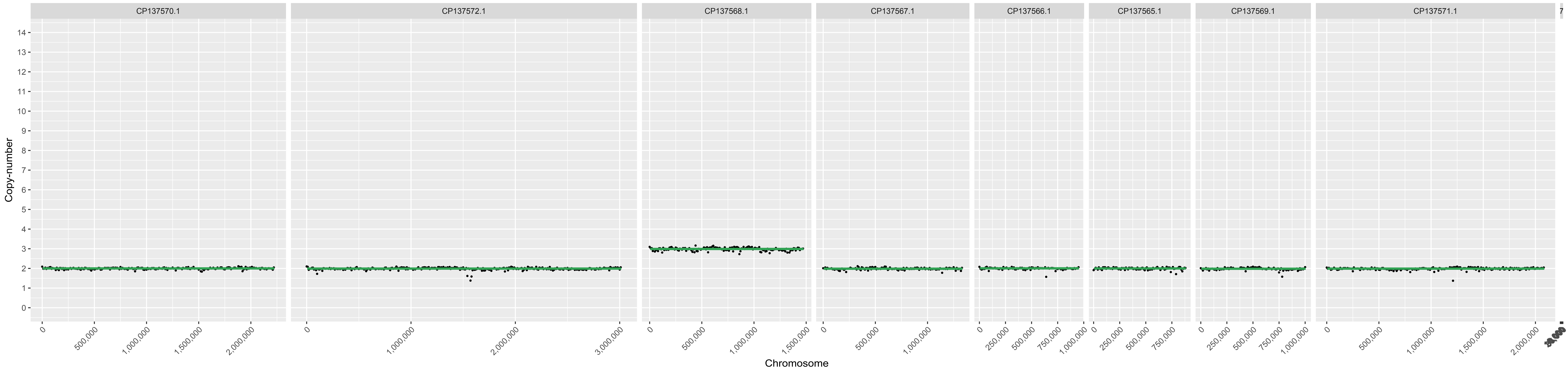

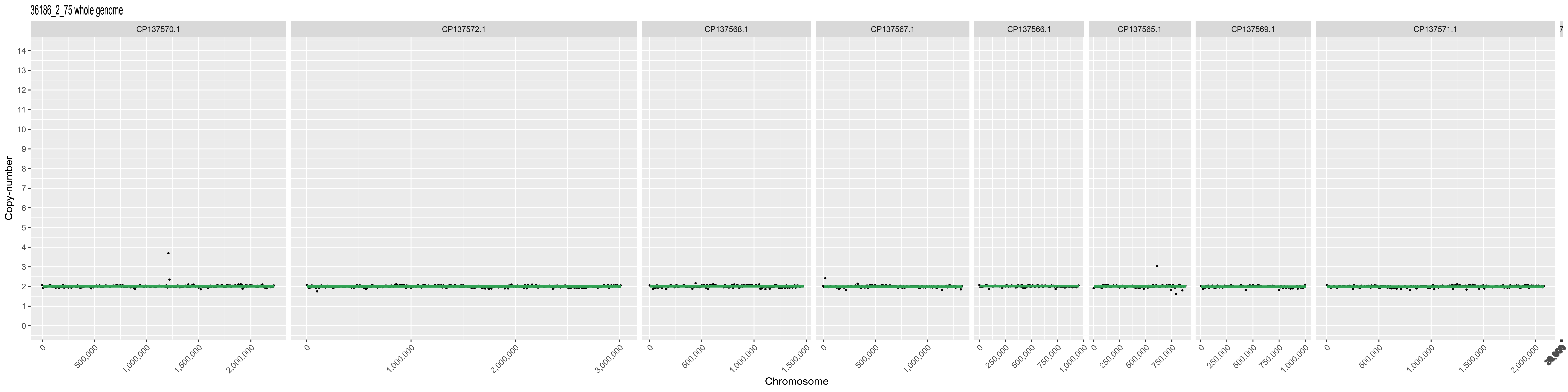
36186_2_75 whole genome
CP137570.1
CP137572.1
CP137568.1
CP137567.1
CP137566.1
CP137565.1
CP137569.1
CP137571.1
Copy-number
0
500,000
1,000,000
1,500,000
2,000,000
0
1,000,000
2,000,000
3,000,000
0
500,000
1,000,000
1,500,000
0
500,000
1,000,000
0
250,000
500,000
750,000
1,000,000
0
250,000
500,000
750,000
0
250,000
500,000
750,000
1,000,000
0
500,000
1,000,000
1,500,000
2,000,000
Chromosome

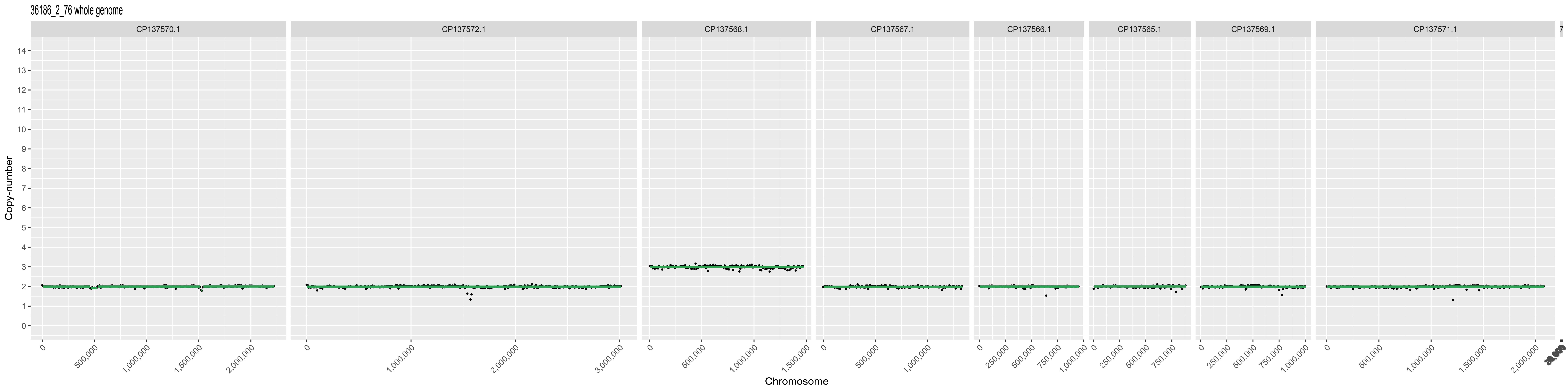
36186_2_76 whole genome
CP137570.1
CP137572.1
CP137568.1
CP137567.1
CP137566.1
CP137565.1
CP137569.1
CP137571.1
Copy-number
Chromosome

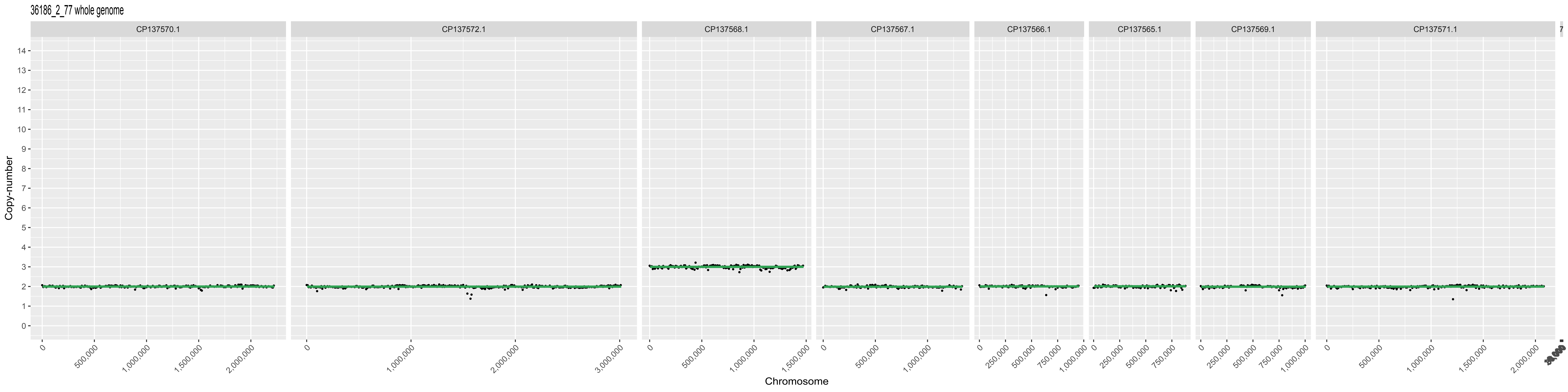

36186_2_77 whole genome
CP137570.1
CP137572.1
CP137568.1
CP137567.1
CP137566.1
CP137565.1
CP137569.1
CP137571.1
Copy-number
Chromosome

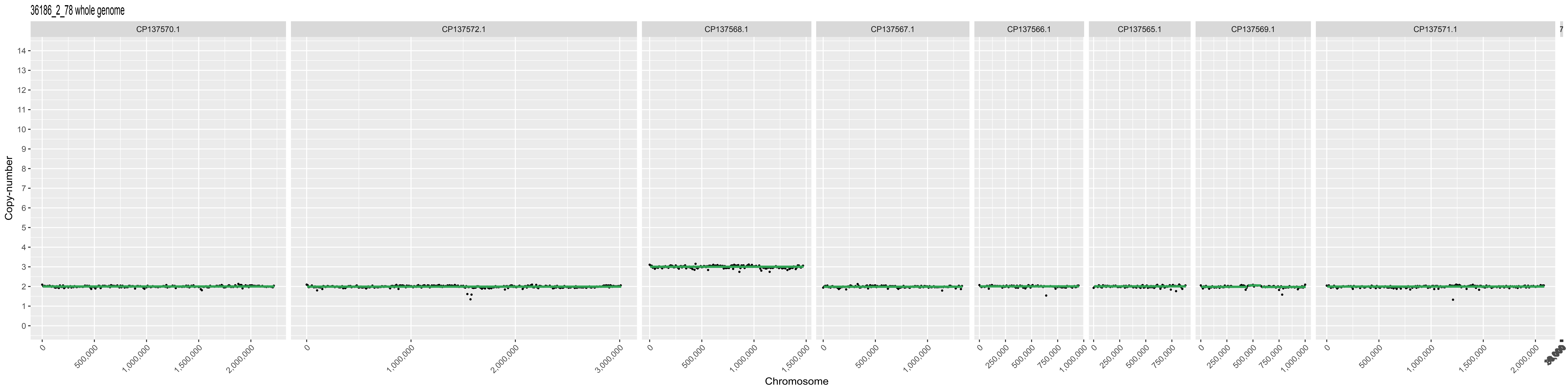

36186_2_78 whole genome
CP137570.1
CP137572.1
CP137568.1
CP137567.1
CP137566.1
CP137565.1
CP137569.1
CP137571.1
Copy-number
Chromosome

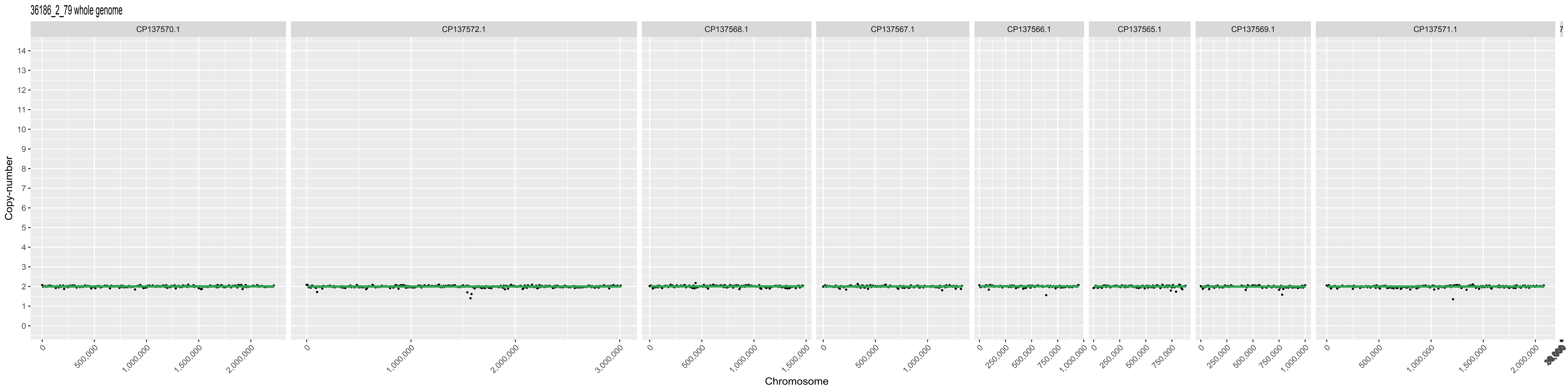
36186_2_79 whole genome
CP137570.1
CP137572.1
CP137568.1
CP137567.1
CP137566.1
CP137565.1
CP137569.1
CP137571.1
Copy-number
Chromosome

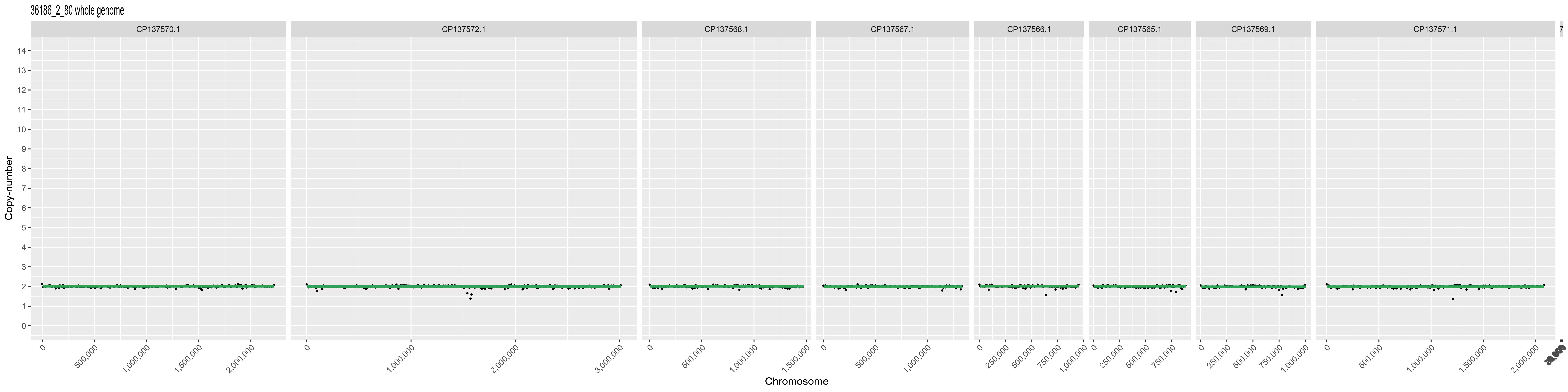
36186_2_80 whole genome
CP137570.1
CP137572.1
CP137568.1
CP137567.1
CP137566.1
CP137565.1
CP137569.1
CP137571.1
Copy-number
Chromosome
500,000
1,000,000
1,500,000
2,000,000
3,000,000
250,000
750,000

36186_2_82 whole genome

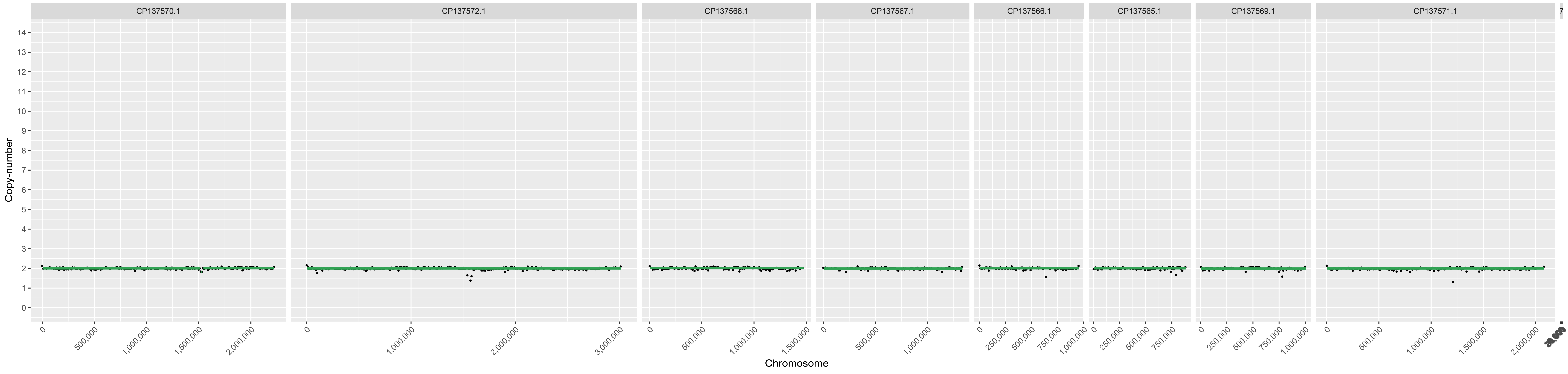

36186_2_83 whole genome

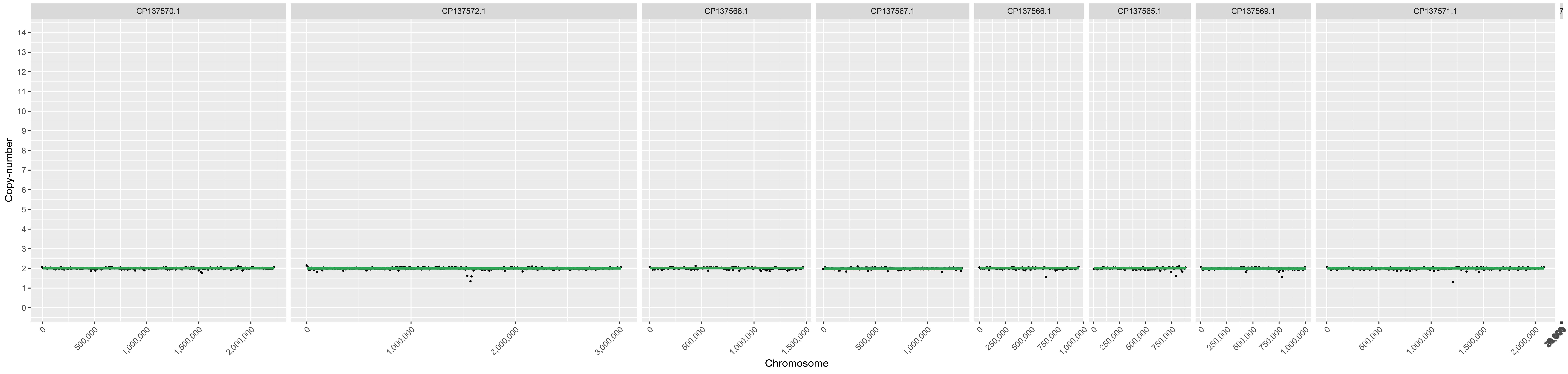

36186_2_85 whole genome

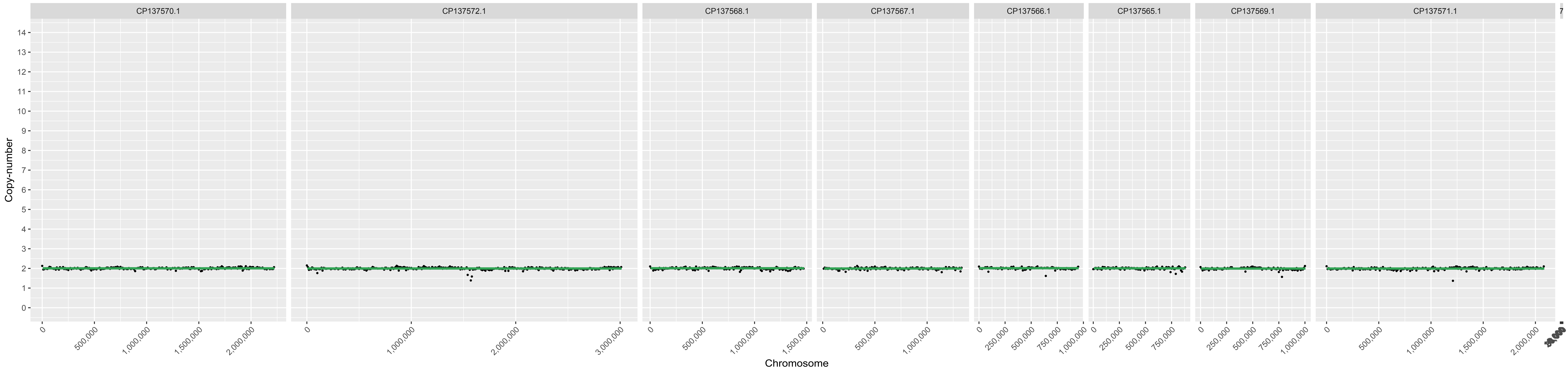

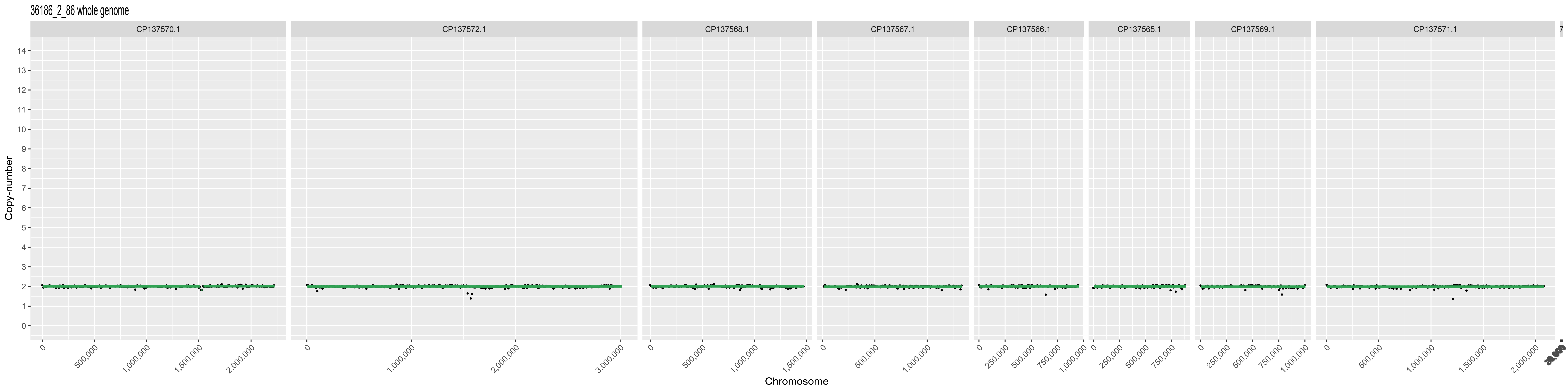

36186_2_86 whole genome
CP137570.1
CP137572.1
CP137568.1
CP137567.1
CP137566.1
CP137565.1
CP137569.1
CP137571.1
Copy-number
Chromosome

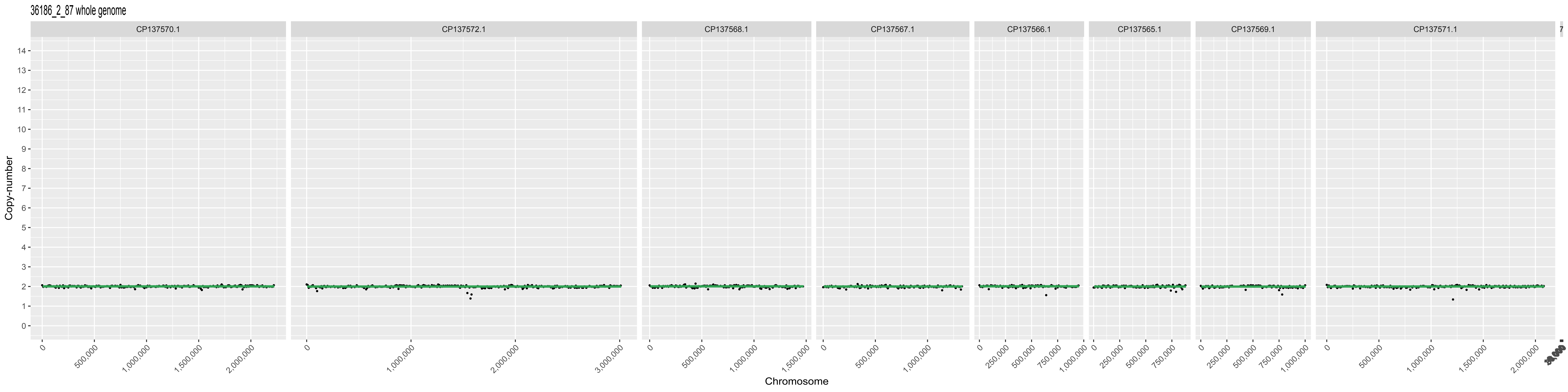

36186_2_87 whole genome
CP137570.1
CP137572.1
CP137568.1
CP137567.1
CP137566.1
CP137565.1
CP137569.1
CP137571.1
Copy-number
Chromosome

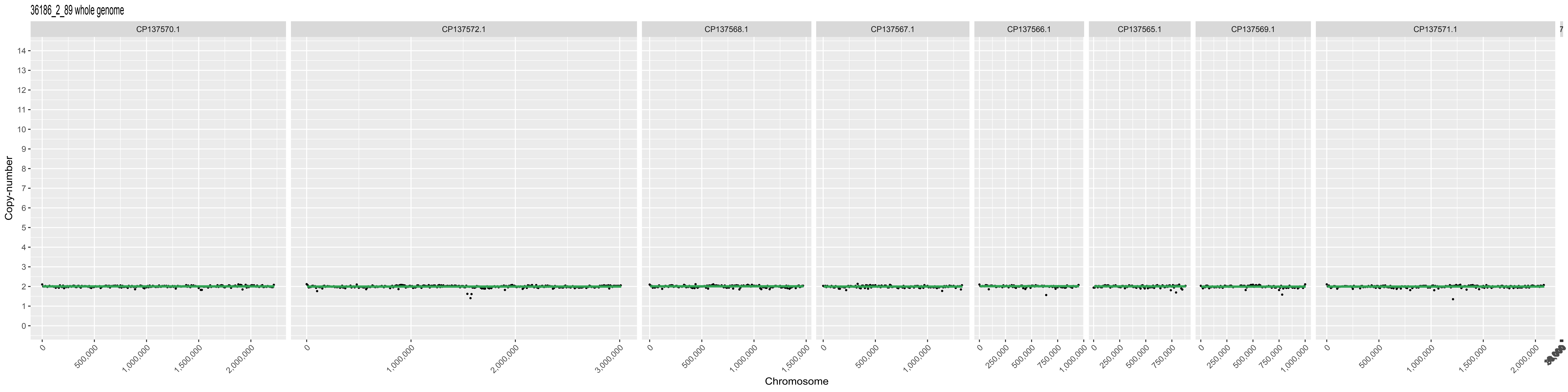
36186_2_89 whole genome
CP137570.1
CP137572.1
CP137568.1
CP137567.1
CP137566.1
CP137565.1
CP137569.1
CP137571.1
Copy-number
Chromosome

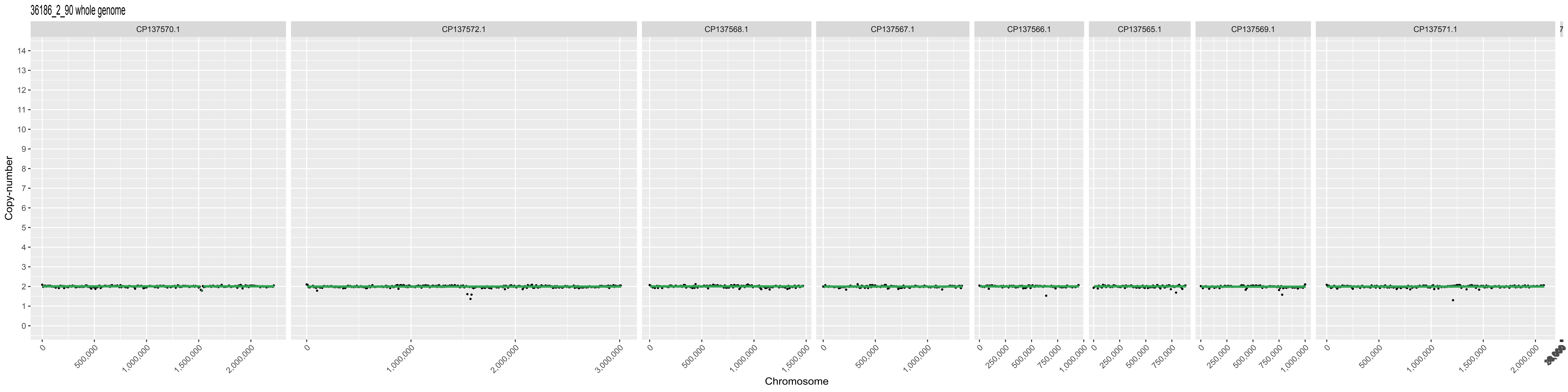

36186_2_90 whole genome
CP137570.1
CP137572.1
CP137568.1
CP137567.1
CP137566.1
CP137565.1
CP137569.1
CP137571.1
Copy-number
Chromosome

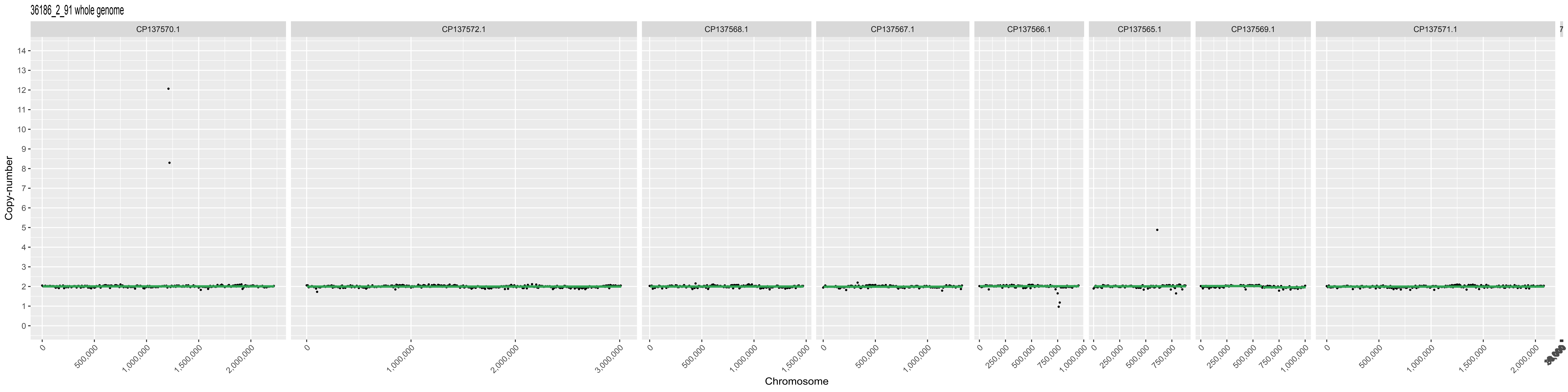
36186_2_91 whole genome
CP137570.1
CP137572.1
CP137568.1
CP137567.1
CP137566.1
CP137565.1
CP137569.1
CP137571.1
Copy-number
Chromosome

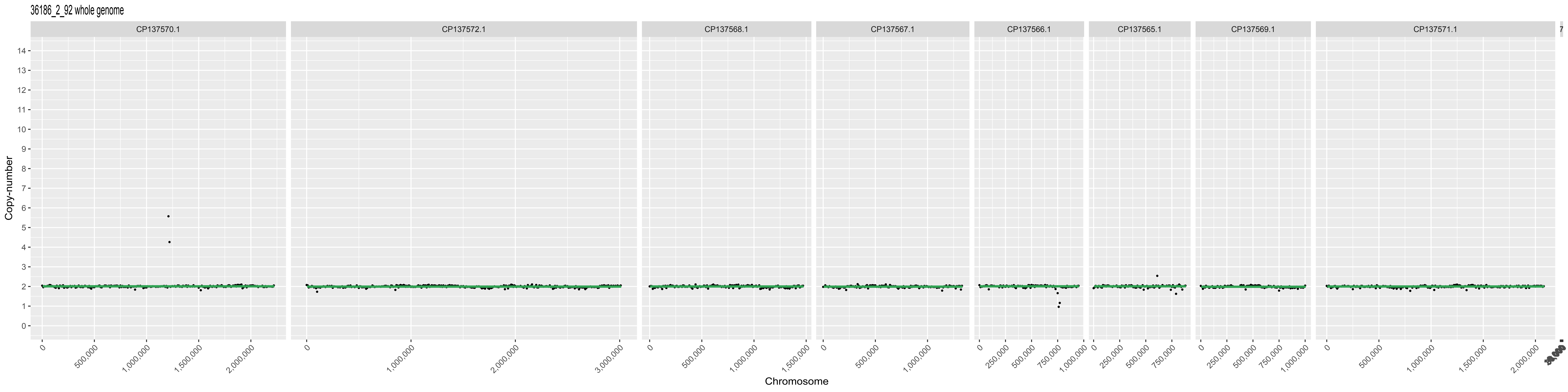

36186_2_92 whole genome
CP137570.1
CP137572.1
CP137568.1
CP137567.1
CP137566.1
CP137565.1
CP137569.1
CP137571.1
Copy-number
Chromosome

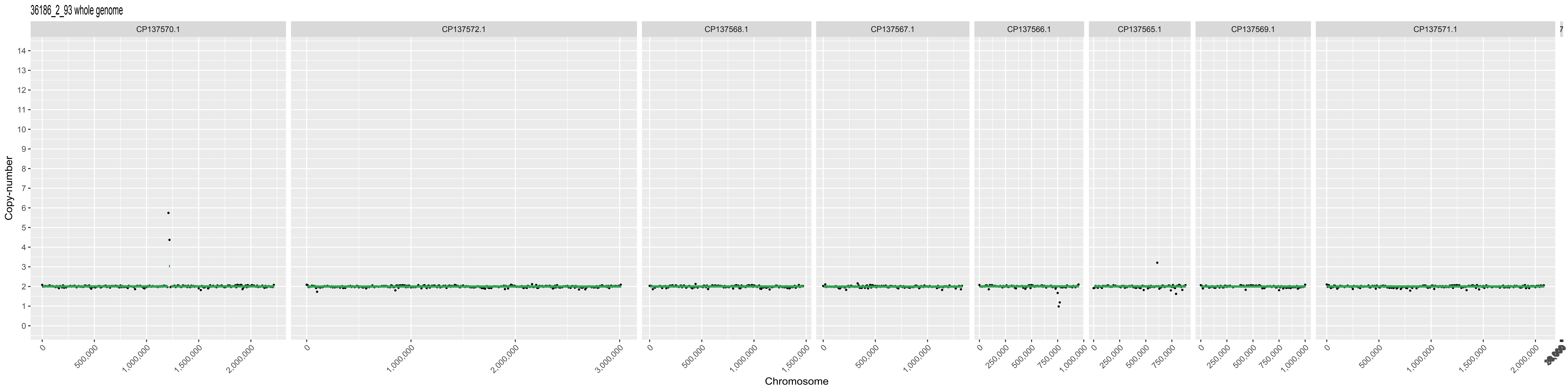

36186_2_93 whole genome
CP137570.1
CP137572.1
CP137568.1
CP137567.1
CP137566.1
CP137565.1
CP137569.1
CP137571.1
Copy-number
Chromosome
0
250,000
500,000
750,000
1,000,000
1,500,000
2,000,000
3,000,000

36186_2_94 whole genome

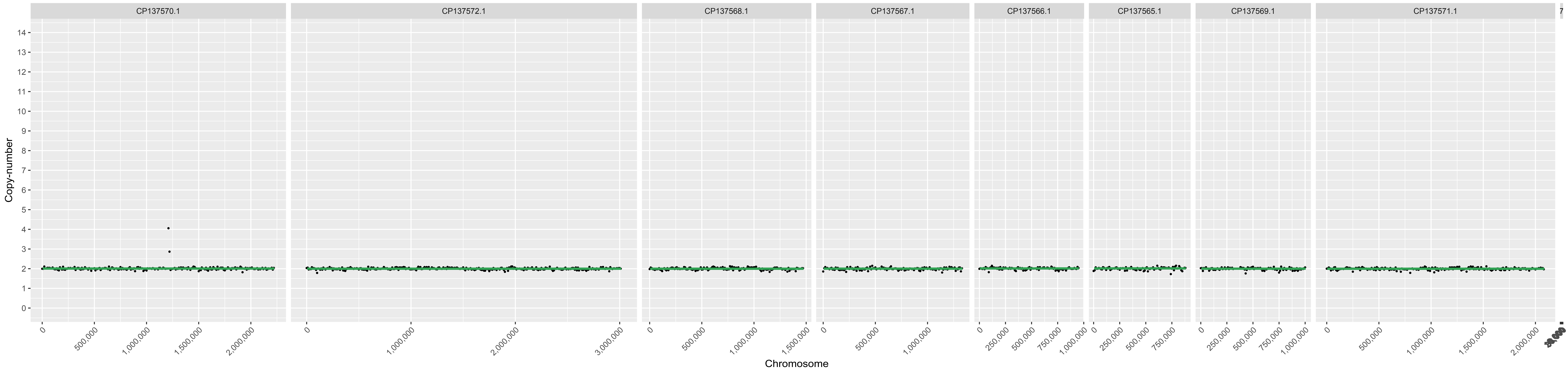

36186_2_95 whole genome

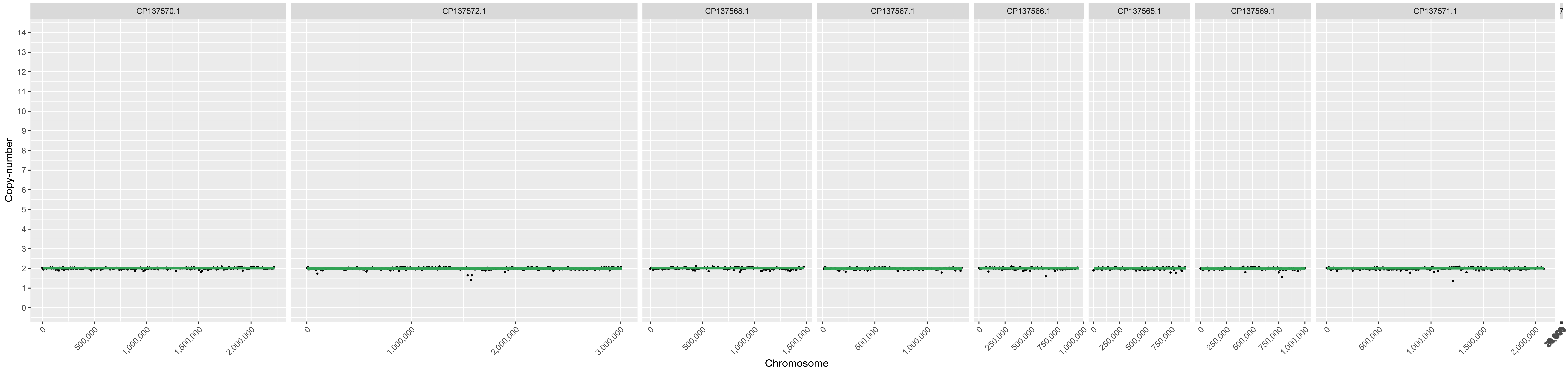

36186_2_96 whole genome

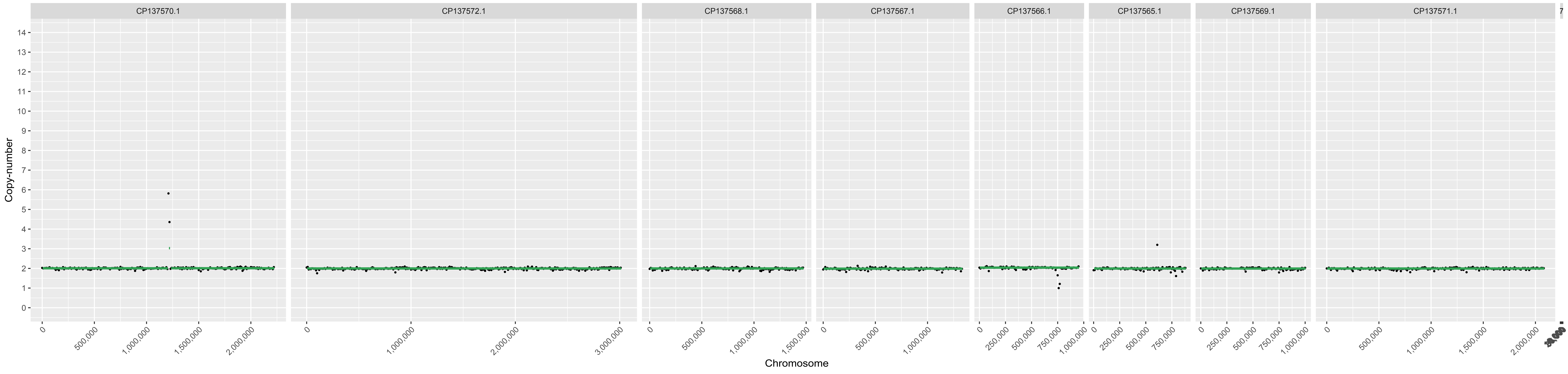
